# YAP signaling regulates the cellular uptake and therapeutic effect of nanoparticles

**DOI:** 10.1101/2023.03.10.532035

**Authors:** Marco Cassani, Soraia Fernandes, Jorge Oliver-De La Cruz, Helena Durikova, Jan Vrbsky, Marek Patočka, Veronika Hegrova, Simon Klimovic, Jan Pribyl, Doriana Debellis, Petr Skladal, Francesca Cavalieri, Frank Caruso, Giancarlo Forte

**Author notes:** Corresponding authors. (GF); (MC). Institute for Bioengineering of Catalonia (IBEC), The Barcelona Institute for Science and 16 Technology (BIST), Barcelona, Spain.

## Abstract

Interactions between living cells and nanoparticles have been extensively studied to enhance the delivery of therapeutics. Nanoparticles size, shape, stiffness and surface charge have been regarded as the main features able to control the fate of cell-nanoparticle interactions. However, the clinical translation of nanotherapies has so far been limited, and there is a need to better understand the biology of cell-nanoparticle interactions. This study investigated the role of cellular mechanosensitive components in cell-nanoparticle interactions. We demonstrate that the genetic and pharmacologic inhibition of yes-associated protein (YAP), a key component of cancer cell mechanosensing apparatus and Hippo pathway effector, improves nanoparticle internalization in triple-negative breast cancer cells regardless of nanoparticle properties or substrate characteristics. This process occurs through YAP-dependent regulation of endocytic pathways, cell mechanics, and membrane organization. Hence, we propose targeting YAP may sensitize triple negative breast cancer cells to chemotherapy and increase the selectivity of nanotherapy.

## Introduction

Yes-Associated Protein (YAP) is a mechanically activated downstream effector of the Hippo pathway (*1*) that plays a critical role in embryogenesis by controlling the size and shape of organs through the proliferation of embryonic parenchymal cells, such as cardiomyocytes and hepatocytes (*2, 3*). Similar to its paralog protein TAZ, which is encoded by the *WWTR1* gene, YAP acts in a pleiotropic fashion by interacting with tissue- and stage-specific transcription factors (*4, 5*), primarily those of the TEAD family (*6-9*). YAP is commonly overexpressed in many solid tumors including breast, lung, colorectal, pancreatic, and liver carcinomas, as well as melanoma and glioma, during their growth, progression and metastasis (*10-13*). YAP has been shown to promote tumor survival by driving tumor immune evasion through the activation of PD-L1 transcription and by rewiring macrophage response to a pro-tumor phenotype (*14*). Additionally, it appears to inhibit autophagy-related cell death (*15*), and drive tumor resistance to targeted therapy and chemotherapy, supposedly through the stimulation of pro-survival and anti-apoptotic genes (*14*). YAP overexpression has been linked to poor prognosis and survival in patients with breast cancer, as well as in other tumour types (*16, 17*).

Our group has recently demonstrated that YAP-mediated activation of cell adhesion genes drives the stiffening of CAL51 triple-negative breast cancer (TNBC) cells (*18*). TNBC defines a subtype of breast cancer characterized by aggressive behavior, frequent relapses, and resistance to chemotherapy (*19, 20*). We have subsequently shown that targeting YAP via mechanical, pharmacological or genetic strategies prevents breast cancer cells from undergoing epithelial-to-mesenchymal transition (EMT) and migration, favoring instead the acquisition of a terminally differentiated phenotype of adipocytes (*21*).

Increased extracellular matrix (ECM) stiffness is a common feature of solid tumors (*22, 23*), and the expression of EMT transition markers is often used to gauge the aggressiveness of breast cancer (*24*). Interestingly, YAP hyperactivation has been recently discovered to play a key role in enabling cancer-associated fibroblasts (CAFs) to induce tumor stroma stiffening and promote malignant cell invasion and metastatization (*25*). Given its dual role in CAFs and tumor cells as both a mechanosensitive protein and a proto-oncogene, YAP is now viewed as a critical component in the positive feedback loop that leads to stroma stiffening and cancer dissemination. Recently, manipulating the mechanical properties of cells or substrates has been proposed as a plausible strategy to control nanoparticle binding and internalization, hence paving the way to using nanoparticles for the *mechanotargeting* of primary or metastatic cancer cells (*26, 27*). Considering this, understanding the interactions between biological systems and nanomaterials has become a major goal of nanomedicine, with the aim of designing nanomaterials that can effectively interact with living cells (*28, 29*).

The design and effectiveness of nanomedicines for cancer therapy depend on various physicochemical properties of the nanocarrier, including their size, shape, stiffness, and surface chemistry (*30, 31*). Much research has focused on optimizing these properties to enhance cell-nanoparticle interaction and improve anti-cancer drug delivery (*32, 33*). Notwithstanding significant advances in understanding bio-nano interactions over the last 20 years, much remains unknown about the exact mechanisms underlying cell-nanoparticle interactions (*34*). Nevertheless, unveiling the processes responsible for these interactions at the molecular level may lead to the development of new strategies for enhancing nanoparticle delivery to specific cells of interest (*35*).

Despite YAP is being proposed as a target for novel treatments (*36, 37*), its response to nanoparticles internalization and its potential role in their trafficking has never been investigated.

In the present study, we used TNBC cells to unveil the role of YAP in cell-nanoparticle interactions and show its potential in regulating nanoparticle internalization via the control of membrane organization and cell mechanics. We demonstrate that YAP is responsible for the transcription of genes regulating cell-matrix interactions, ECM deposition, and endocytic pathways in breast cancer cells, and show that its inhibition boosts the internalization of nanoparticles. Finally, we delineate how the intracellular delivery of doxorubicin-loaded liposomes can be enhanced by pharmacologically tampering with YAP activity. In conclusion, by identifying Hippo effector as a determinant of cell-nanoparticle interaction, we propose its inhibition as a viable therapeutic strategy for improving nanodrug delivery to triple negative breast cancer cells.

## Results

### YAP depletion affects the mechanical, physical properties and the membrane organization of CAL51 TNBC cells

We first investigated the effect of YAP depletion on the morphology of CAL51 TNBC cells, which are characterized by constitutively high levels of YAP expression and activity (*21*). Using CRISPR/CAS9 technology, we generated a stable YAP-deficient mutant CAL51 cell line (described in (*18*)), and confirmed YAP depletion through confocal microscopy (Fig. 1A) and western blot analyses (Fig. 1B). Next, we used atomic force microscopy (AFM) to determine the effect of YAP depletion on the mechanical properties of CAL51 cells and found a significant reduction in Young’s modulus in YAP -/- cells (Fig. 1C). Moreover, compared to CAL51 WT cells, YAP -/- cells displayed reduced surface area, both as determined by actin coverage and membrane extension (Fig. 1D-E), as well as perturbed cell morphology, resulting from their inability to spread over the adhesion surface (Fig. 1F; see fig. S1). These changes were due to the failure of the mutant cells to assemble focal adhesions and form proper cytoskeleton (Fig. 1G) (*18, 38*). Importantly, YAP depletion did not affect the proliferation and viability of CAL51 TNBC cells (see fig. S2).

**Fig. 1.**
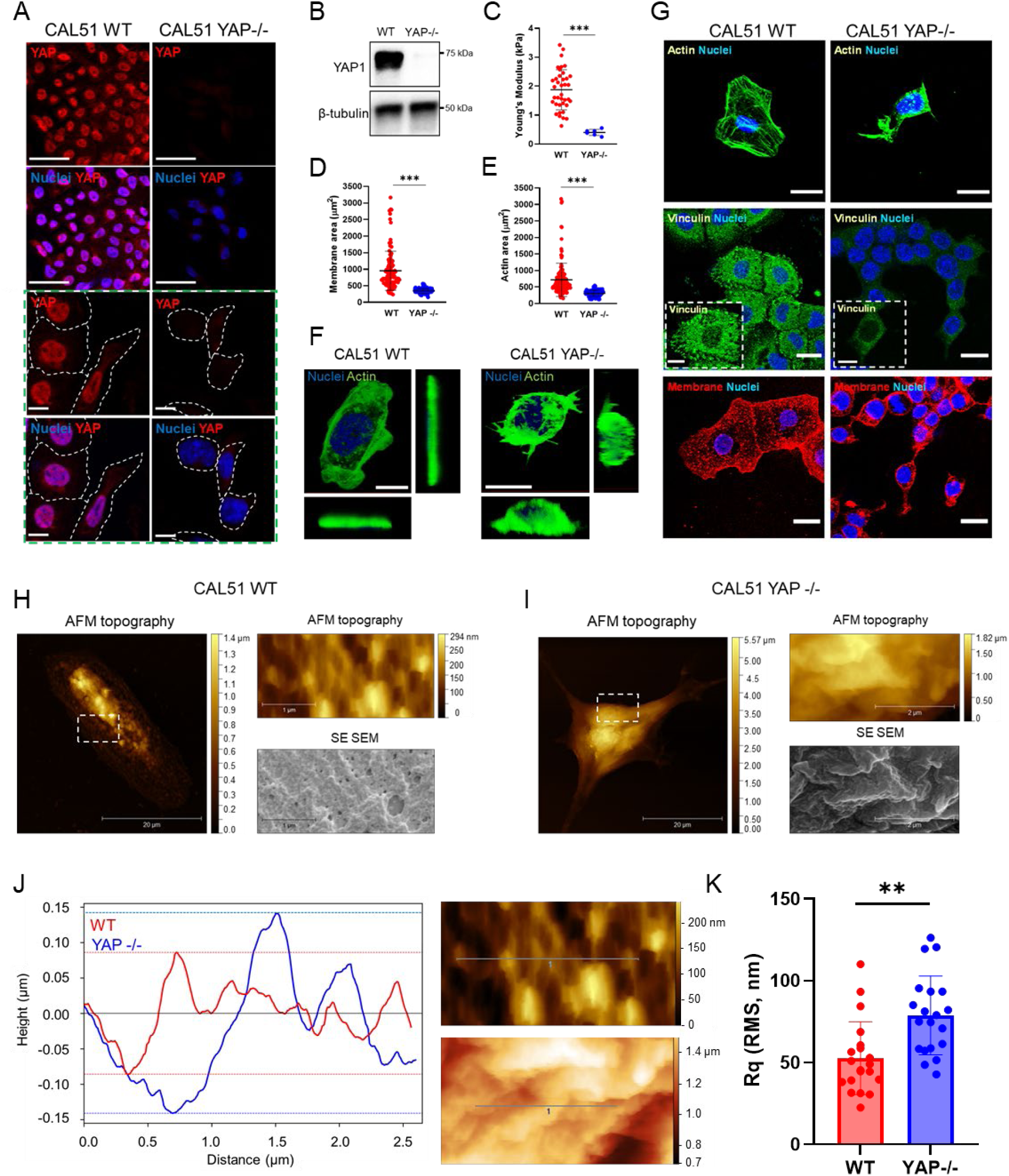
YAP depletion affects CAL51 adhesion, mechanics, morphology, and membrane properties. A) Representative confocal images depicting YAP expression in WT or YAP -/- CAL51 cells. Cells were stained for YAP (AF488), and nuclei were counterstained with DAPI. Scale bar: 50 μm. The green dashed line box shows higher magnification pictures. Scale bar: 10 μm. B) Western blot analysis showing the levels of YAP protein in WT or YAP -/- CAL51 cells. β-tubulin was used for protein loading normalization. C) Dot plot representation of the Young’s modulus analysis of WT or YAP -/- CAL51 cells as measured by atomic force microscopy (AFM). WT CAL51: n = 80; YAP -/- CAL51: n = 10. ***p < 0.001. D) Dot plot analysis of WT or YAP -/- CAL51 total membrane area. Cells were stained with Alexa Fluor 488-labelled wheat germ agglutinin (WGA-488, green). n > 100 cells. ***p < 0.001. E) Dot plot analysis of WT or YAP -/- CAL51 cell surface area calculated based on the total actin coverage of the cells. Cells were stained with Alexa Fluor 488-labelled Phalloidin (Pha-488, green). n > 100. ***p < 0.001. F) CAL51 WT (left) and YAP -/- cells (right) 3D reconstruction. Cells were stained with DAPI and WGA-488 (green). Scale bar: 20 μm. G) Representative confocal images of WT or YAP -/- CAL51 cells stained for DAPI and actin (Pha-488, green, top), vinculin (AF488, green, middle), and membrane (WGA-647, red, bottom), respectively. Scale bar: 20 μm. The insets display high magnification images. Scale bar: 10 μm. Correlative Probe and Electron Microscopy (CPEM) imaging of CAL51 WT (H) and CAL51 YAP -/- (I) cells. AFM and SEM images are shown. White dashed line boxes indicate the detail of the magnifications shown as AFM and SEM images on the right of each main micrograph. J) Plot displaying the profile of the membrane roughness as determined for WT (red) and YAP -/- (blue) CAL51 cells in the region highlighted in the SEM images on the right (red dashed line, top for CAL51 WT; blue dashed line, bottom for YAP -/- CAL51). The roughness profile was calculated on the deconvolved images. Scale bar: 0.5 μm. K) Mean square roughness of the height irregularity (Rq) measured on WT (red) and YAP -/- (blue) CAL51cells. n = 20. Statistical analysis was performed by unpaired t-test using Welch’s correction; **p < 0.01.

Given the striking change in morphology and mechanics exhibited by YAP -/- CAL51 cells compared to the wild type control, we analyzed their membrane structure using Correlative Probe and Electron Microscopy (CPEM) by LiteScope™. This technique combines AFM and scanning electron microscope (SEM) to characterize 3D surface in situ, estimate surface roughness, and perform height/depth profiling with precise AFM tip navigation (*39*). The analysis demonstrated that YAP depletion and the following decrease in membrane tension led to the remodeling of the plasma membrane in CAL51 TNBC cells and the emergence of dynamic features connected with extensive cellular reorganization (Fig. 1H-I). Previous literature has reported the appearance of various membrane structures including blebs and vacuole-like dilations upon the reduction of cell strain (*40*). In YAP -/- cells, we observed the emergence of ripple-like deformations (Fig. 1J), which increased the roughness of the cell membrane, as quantified through height irregularity (Rq, Fig. 1K).

### YAP transcriptional activity controls the expression of genes involved in membrane organization and endocytosis in CAL51 TNBC cells

Given the substantial effects that YAP depletion played on the structure of cell membrane, we hypothesized that the changes in the membrane of YAP -/- cells might be reflected in their transcriptional landscape. We adopted RNA sequencing (RNA-seq) to investigate the regulation of genes encoding for proteins involved in plasma membrane organization. Overall, RNA-seq revealed a total of 4,219 differentially expressed genes in YAP -/- cells compared to WT CAL51 cells, with 1,925 of them being downregulated and 2,294 upregulated following YAP depletion (see fig. S3a-c). The most represented gene ontology (GO) annotations were connected to ECM organization, cell migration, and cell adhesion in WT cells (see fig. S3d), while genes related to integral components of the plasma membrane were found among the cellular components in YAP -/- counterpart (see fig. S3e).

In line with our findings regarding cell membrane organization (Figures 1H-M), we detected the presence of amphiphysin I (*AMPH1*) among the genes being affected the most by YAP depletion. *AMPH1* encodes for a protein that senses and generates membrane curvatures, is implicated in clathrin-mediated endocytosis (*41*) and was significantly upregulated in CAL51 YAP -/- cells (log2Fc 7.69, P < 0.05, Fig. 2A). Together with *AMPH1*, several genes contributing to plasma membrane assembly that were overexpressed in WT CAL51 may explain the differences in cell membrane organization (Fig. 2A). These include *RFTN1* (log2Fc 2.79), *EHD2* (log2Fc 2.4), *MYOC* (log2Fc 5.83) and *PITPNM1* (log2Fc 2.86) (*42-45*).

**Fig. 2.**
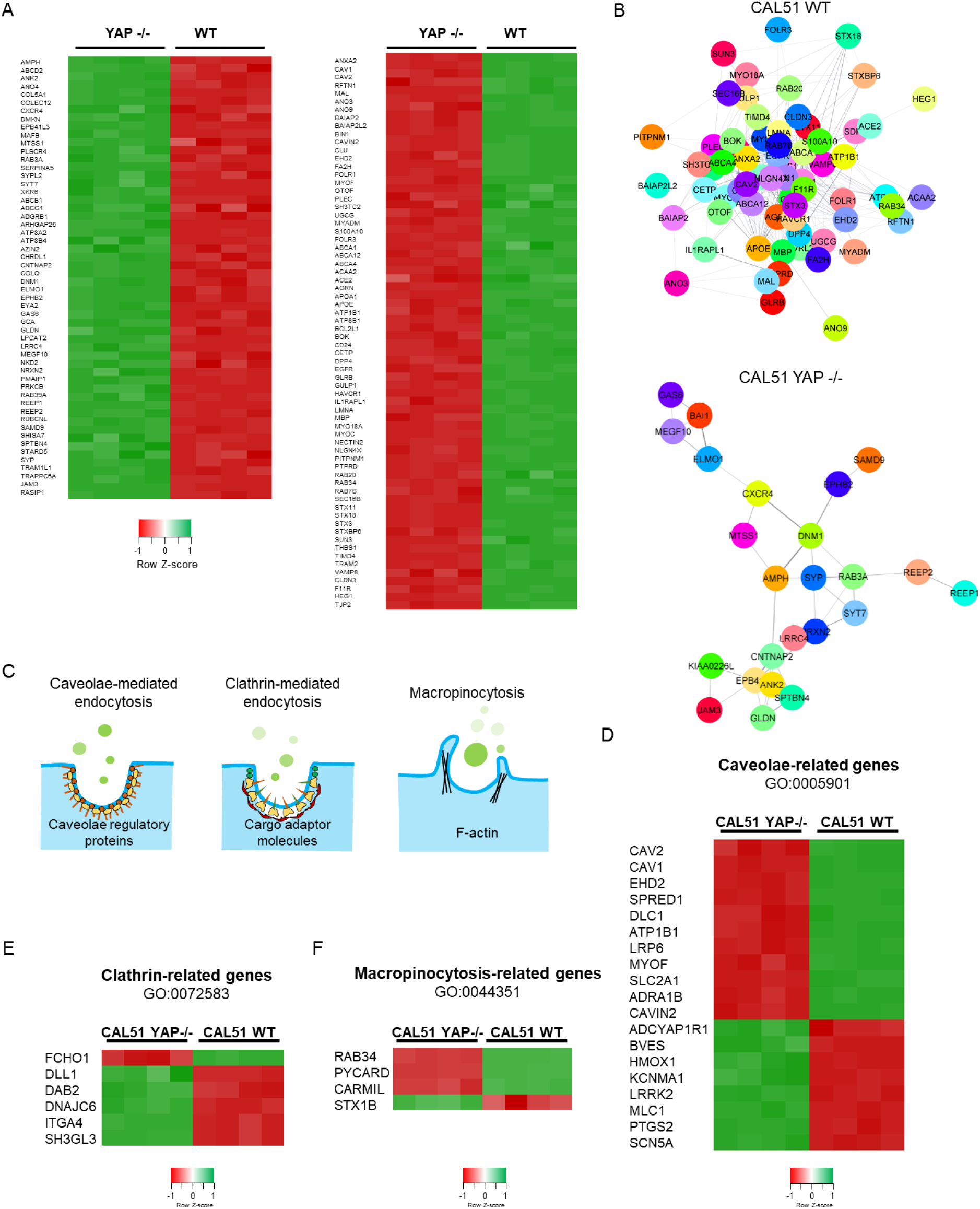
YAP depletion alters the expression of genes related to membrane organization and endocytosis pathways. A) Heatmap of the relative expression of significantly regulated genes associated with the membrane organization network in YAP -/- and WT CAL51 cells. n = 4 (P adj < 0.05, log2Fc > ǀ2ǀ). B) STRING PPI network of differently expressed proteins involved in membrane organization in WT (top) and YAP -/- CAL51 (bottom) cells obtained from Cytoscape (P adj < 0.05, log2Fc > ǀ2ǀ, confidence cutoff 0.4). C) Graphical representation of the main mechanisms of endocytosis, i.e. caveolae-related endocytosis, clathrin-related endocytosis and macropinocytosis, investigated in the present study. D) heatmap of genes involved in endocytosis pathways for caveolae-related genes (D). E) heatmap of genes involved in clathrin-mediated endocytosis pathways. (F). Heatmap of genes involved in macropinocytosis (P adj < 0.05, log2Fc > ǀ1ǀ).

GO enrichment analysis identified several molecular functions being differentially regulated that were connected to plasma membrane network components, with high scores for lipid raft organization, localization, and assembly (see fig. S4a). Interestingly, lipid rafts, responsible for membrane heterogeneity, strongly impacts the stability and functionality of the plasma membrane at the nanoscale, thus directly supporting high sub-compartmentalization and intrinsic organization (*46*). In addition, STRING PPI analysis yielded a highly clustered network containing 68 nodes and 384 edges for WT CAL51, while 54 nodes and 39 edges were identified for YAP-depleted cells (Fig. 2B). WT CAL51 cells showed a significantly higher number of interactors and interconnections between the key elements of the membrane organization network compared to YAP -/- cells, possibly indicating a higher degree of membrane complexity. The cluster coefficients for both conditions were quite similar (0.394 for WT and 0.5 for YAP -/-). More importantly, RNA-seq analysis revealed the differential expression of genes encoding proteins involved in endocytosis (Fig. 2C; see fig. S4b), a process that appears to be dysregulated in tumors (*47*). Therefore, the RNA-seq analysis was extended to examine the expression of transcripts involved in the trafficking of intracellular organelles and endocytic pathways in YAP -/- cells compared to CAL51 WT cells. This comparison led to the identification of differentially expressed genes involved in caveolae-mediated endocytosis (Fig. 2D), clathrin-mediated endocytosis (Fig. 2E), and macropinocytosis (Fig. 2F). While the number of differentially regulated genes in the caveolae-mediated pathway was similar between WT and YAP -/- cells (11 in WT and 8 in YAP -/-, although YAP -/- cells displayed overall higher upregulation; see fig. S4b), clathrin-related and macropinocytosis-related genes showed the most striking difference, with more genes of the clathrin-mediated pathway upregulated in YAP -/- cells (5 genes in YAP -/- vs. 1 gene in WT) and more genes of the macropinocytosis-mediated pathway upregulated in WT cells (1 gene in YAP -/- vs. 3 genes in WT).

To further corroborate these findings, we performed a GO enrichment analysis on the genes involved in endocytosis that were differentially expressed in WT and YAP -/- cells. The analysis revealed significant differences between WT and YAP -/- cells in annotations for the negative and positive regulators of endocytosis (see fig. S5 and S6). For the negative regulators (GO:0045806), WT cells showed high scores for the regulation of endocytic vesicles (*P* = 0.001) and early endosomes (*P* = 0.002) compared to YAP -/- cells (*P* = 0.13 for endocytic vesicles, *P* = 0.03 for early endosomes), whereas YAP -/- cells yielded high scores for endosomal recycling genes (*P* = 0.014). These findings indicate that YAP depletion in triple negative breast cancer cells lead to the downregulation of transcripts encoding proteins controlling the endocytic pathways (see fig. S5). The positive regulators (GO:0045807), on the other hand, were more abundant in the absence of YAP: relative to WT CAL51, YAP -/- cells showed significant scores for the terms *positive regulators of the endocytic vesicle* (*P* = 3.11×10^-5^ in YAP -/- vs. *P* = 0.006 in WT) and *cytoplasmic vesicle membrane* (*P* = 0.000476 in YAP -/- vs. *P* = 0.02 in WT), while CAL51 WT had high scores for *collagen-containing extracellular matrix* (*P* = 7.56×10^-5^) and *secretory granule lumen* (*P* = 0.00035; see fig. S6). Taken together, these results indicate that YAP depletion in CAL51 TNBC cells determines an alteration in the genes involved in endocytosis.

### YAP regulates the entry of nanoparticles in CAL51 TNBC cells

Encouraged by these findings, we hypothesized that YAP might play a role in the internalization of nanoparticles and nanodrugs. We figured this might explain how Hippo pathway effector promotes TNBC resistance to chemotherapy (*48*). To test this hypothesis, we treated WT and CAL51 YAP -/- cells with inert polystyrene (PS) nanoparticles (NPs), which have been widely used in bio-nano interaction studies due to their tunable size and ease of functionalization (*49, 50*). Carboxylated PS nanoparticles with 200 and 900 nm diameter (PS200 and PS900) were first labelled with carboxytetramethylrhodamine using EDC chemistry and characterized *via* transmission electron microscopy (TEM), spectrofluorometer, and dynamic light scattering (DLS) (see fig. S7a-e). Next, WT and YAP -/- CAL51 cells were incubated with PS200 and PS900, and nanoparticle binding to the cells evaluated using flow cytometry and confocal microscopy. Our flow cytometry results showed that already after 4 h incubation, PS nanoparticles could be found preferentially bound to YAP -/- CAL51 cells as compared to their WT counterparts (Fig. 3A; see fig S8 and S9a). This effect was independent of the nanoparticle size and could be further confirmed by confocal analysis. Through this analysis we – in fact – showed that a higher number of nanoparticles co-localized with the membrane of YAP -/- compared to WT cells (Fig. 3B-C; see fig. S9b). Interestingly, CPEM analysis allowed us to visualize the detailed morphology of the nanoparticles in contact with CAL51 cells, further corroborating these findings. In WT cells, NPs were bound to the external face of the membrane but not yet internalized after 4 hours (see fig. S10a). They in fact appeared bright on SEM and AFM imaging in CAL51 WT and showed limited co-localization with the cell membrane (see the height profile of the area at the nanoparticle-membrane binding site in fig. S10b). On the contrary, at the same time-point NPs were already surrounded by an organic coating attributable to plasma membrane in cells in which YAP had been depleted (see fig. S10c). This result indicated that the nanoparticles were embedded in the cell membrane, as indicated by the reduction of the height profile at the nanoparticle-membrane binding site (see fig. S10d).

**Fig. 3.**
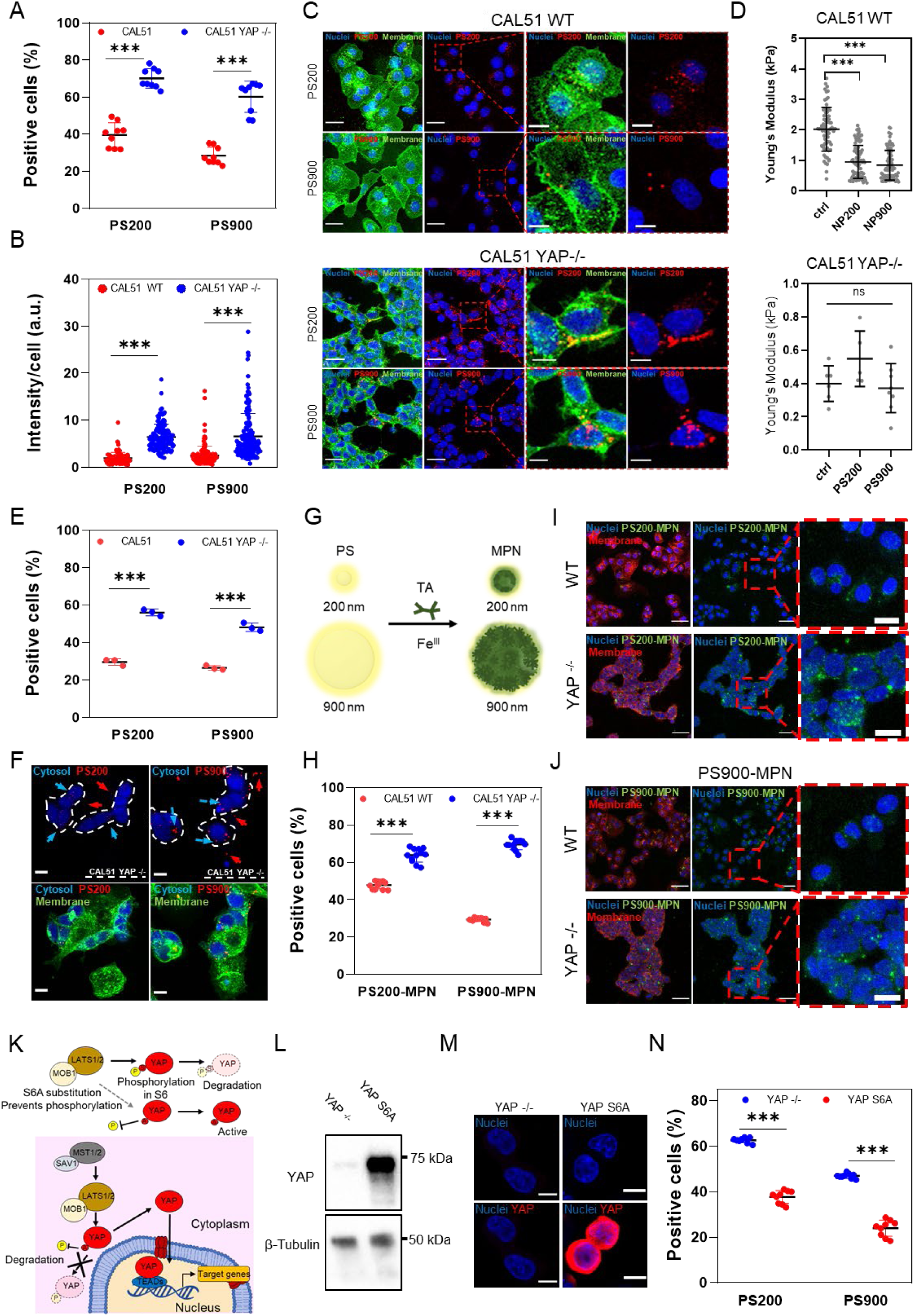
YAP knockout promotes nanoparticle binding and internalization in CAL51 TNBC cells. A) 4-hour cellular uptake of PS200 and PS900 in WT (red) and YAP -/- CAL51 (blue). Two-way ANOVA with Sidak’s correction for multiple comparisons. n = 3; ***p < 0.001. B) Nanoparticle intensity per cell after 4 hours of incubation of CAL51 WT and YAP -/- cells with PS200 and PS900. Statistical analysis was performed using the two-way ANOVA with Sidak’s correction for multiple comparisons. n = 3; *** indicates p < 0.001. n > 100 cells. C) Confocal images of WT (top) and YAP -/- (bottom) CAL51 cells (right) after 4 hours of incubation with PS200 and PS900. Cells were stained with WGA-488 (green) and/or DAPI (blue). Magnified images are displayed inside the red dashed line boxes for each cell and particle type. Scale bar: 25 and 10 μm. D) Young’s modulus analysis of WT (top) and YAP -/- (bottom) cells after 4 hours of incubation with PS200 and PS900, as measured by atomic force microscopy (AFM). Kruskal-Wallis one-way ANOVA. WT CAL51: n = 80; YAP -/- CAL51: n = 10; ***p < 0.001; ns, non-significant. E) 4-hour cellular uptake of PS200 and PS900 in a co-culture of WT (red) and YAP -/- (blue) CAL51. Two-way ANOVA with Sidak’s correction for multiple comparisons. n = 3; ***p < 0.001. F) Confocal images of WT and YAP -/- cells co-culture in the presence of PS200 and PS900 for 4 hours. White dashed line indicates CAL51 YAP -/- cells. Blue arrows indicate particles co-localized with YAP -/- cells, while red arrows indicate particles in contact with WT cells. CAL51 YAP -/- cells are stained with 7-amino-4-chloromethylcoumarin (blue) and whole cell population with WGA-488 (green). Scale bar: 10 μm. G) The surface properties of PS200 and PS900 were modified by coating them with MPN using tannic acid and FeCl_3_. H) 4-hour cellular uptake of PS200-MPN and PS900-MPN for WT (red) and YAP -/- (blue) CAL51. Two-way ANOVA with Sidak’s correction for multiple comparisons. n = 3; ***p < 0.001. I) and J) Confocal images of WT (top) and YAP -/- (bottom) CAL51 cells after 4 hours of incubation with PS200 (I) and PS900 (J). Cells are stained with WGA-647 (red) and DAPI (blue). The particles are displayed in green. Magnified images are displayed inside red dashed line boxes for each cell and particle type. Scale bar: 50 and 10 μm. K) Schematic representation of the Hippo pathway in repressing YAP translocation to the nucleus. Due to a substitution of serine with alanine in six different positions, YAP-S6A cannot be phosphorylated, thus is constitutively active in the cell nucleus. L) Western blot analysis of the levels of YAP protein in CAL51 YAP -/- and in cells transfected with a plasmid carrying a copy of the YAPS6A gene. β-tubulin was used for protein loading normalization M) Confocal images of YAP -/- (CTRL) and YAPS6A CAL51. Cells were decorated with DAPI (blue) and YAP (red). Scale bar: 10 μm. N) 4-hour cellular uptake of PS200 and PS900 in YAP -/- (blue) and YAPS6A (red) CAL51. Two-way ANOVA with Sidak’s correction for multiple comparisons. n = 3; ***p < 0.001.

We next focused on investigating whether the striking difference in particle interaction with cell membrane could be due to the changes in membrane curvature, actin dynamics, and cell mechanosensing as induced by YAP activity (*18, 51*). Hence, we reduced the nanoparticle dosage and increased the incubation time and found enhanced nanoparticle internalization over time in YAP -/- CAL51, regardless of dose and time (see S11a). No significant change was observed in YAP localization in WT cells after nanoparticle binding, suggesting no direct impact on intracellular protein shuttling (see fig. S11b). However, a marked decrease in membrane stiffness was noted in WT CAL51 after 4 h of incubation with nanoparticles, but no such change was seen in YAP-depleted cells (Fig. 3D). Although the effect of nanoparticles on the cell membrane properties is debated, with responses possibly dependent not only on the nanoparticle size but also their composition (*52*), our data suggest that the reduction in membrane rigidity in WT CAL51 did not enhance nanoparticle internalization over time, as YAP -/- cells exhibited higher binding and internalization in all conditions tested. This was likely due to a stronger impact of YAP depletion on cell membrane stiffness than the physical effects of nanomaterial interactions alone. Furthermore, z-stack confocal images and TEM micrographs confirmed the internalization of particles in both WT and YAP -/- cells (see fig. S12a-b), although a higher number of particles was found to bind and co-localize with the cell membrane in the absence of YAP.

Next, we treated WT and CAL51 YAP -/- cells with inhibitors selective for each type of endocytosis to study the impact of YAP on the route of nanoparticle internalization in CAL51 TNBC cells. In particular, the cells were pre-treated for 2 h with cytochalasin D, chlorpromazine, and nystatin to inhibit macropinocytosis, caveolin-mediated, and clathrin-mediated endocytosis, respectively (*33*). The cells were then incubated with PS200 and PS900 for 4 h. The live/dead assay was used to confirm no significant effect on cell viability under the chosen treatment conditions (see fig. S13). The obtained results showed that PS200 primarily entered the cells through clathrin-mediated endocytosis, while PS900 primarily did so through macropinocytosis (see fig. S14a-b), which was in line with previous findings for similar-sized nanoparticles (*53*). After internalization, the particles followed classical endocytic pathways and accumulated in lysosomes within 8 h of incubation (see fig. S14c). These endocytic processes were not affected by the absence of YAP, suggesting that the protein *per se* does not impact endocytosis pathways but rather affects the dynamics of cell-nanoparticle association and internalization.

To evaluate the potential of manipulating tumor cell mechanosensing to enhance nanoparticle delivery in a heterogeneous and complex milieu, we co-cultured WT CAL51 cells with YAP -/- cells in a 1:1 ratio. The latter had been previously labeled with 7-amino-4-chloromethylcoumarin (CellTracker™). The co-culture was incubated with either PS200 or PS900 for 4 h. In line with our cell morphology data (Figure 1), YAP -/- cells had a lower total surface area compared to the parental line due to their tendency to spread less on the substrate (see fig. S15a-b). Interestingly, despite the lower area exposed, YAP-depleted cells bound and internalized a significantly higher number of nanoparticles, as revealed by flow cytometry and confocal analysis (Fig. 3E-F; see fig. S15c). Indeed, z-projection confocal images showed consistent preferential co-localization of nanoparticles within the membrane of YAP -/- CAL51 cells (see fig. S16a-b). This result contradicts previous studies linking a higher cell surface area with a higher internalization rate (*52, 54*), and suggests that YAP mechanosensing and cell mechanics outplay cell surface area in nanoparticle binding.

Next, we tested whether the internalization of nanoparticles was affected by their surface coating and charge. These parameters are crucial in bio-nano interaction studies and have been extensively studied to optimize nanoparticle design for efficient targeting or escape from specific cell types for improved therapy delivery (*55*). To determine the impact of nanoparticle surface properties on the differential internalization rate seen in CAL51 cells with or without YAP, PS nanoparticles were coated with the metal phenolic network (MPN) to alter their physicochemical properties, such as surface charge and free energy (*56*). Briefly, the MPN coating was applied to the surface of 5-((5-Aminopentyl)thioureidyl)fluorescein-labeled PS200 and PS900 nanoparticles by the assembly of tannic acid (TA) and FeCl_3_ according to a previously described protocol (*50*). The application of the MPN coating formed PS200-MPN and PS900-MPN (Fig. 3G; see fig. S17a), and the success of the coating was confirmed using a spectrophotometer and dynamic light scattering (DLS) analysis (see fig. S17b-e). To examine cell-nanoparticle interactions, WT and YAP -/- CAL51 cells were incubated with PS200-MPN and PS900-MPN for 4 h. Flow cytometry and confocal analysis revealed higher binding and internalization in YAP -/- cells compared to WT cells, supporting the previous results observed using non-coated PS200 and PS900 nanoparticles (Fig. 4H-J; see fig. S18a-b).

**Fig. 4.**
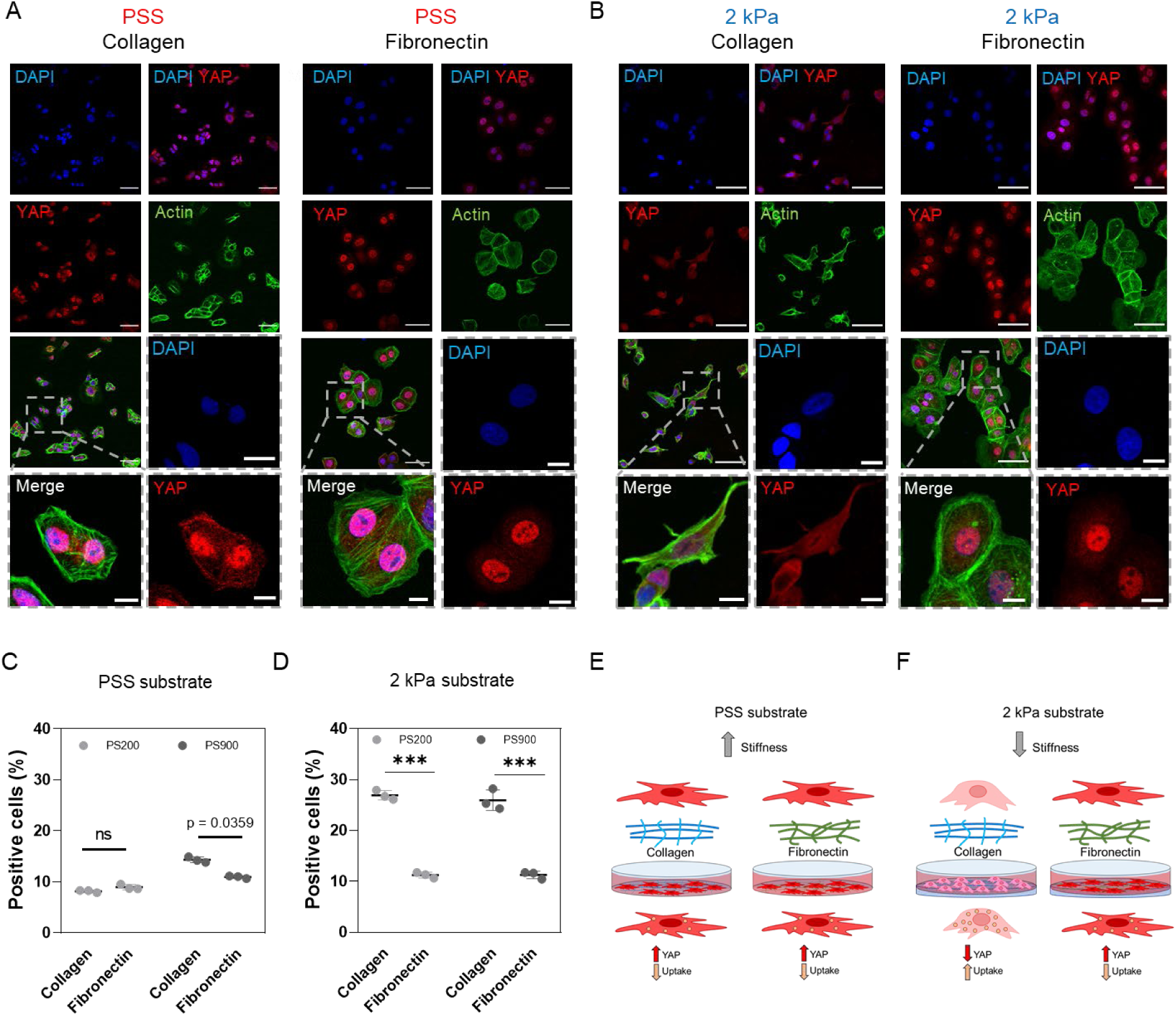
Substrate mechanics impairs nanoparticle uptake through YAP. A) Confocal images of CAL51 WT cells grown on a stiff polystyrene substrate coated with collagen (left) or fibronectin (right). Cells were stained with DAPI (blue), Pha-488 (green), and YAP (red). Magnified images are displayed inside gray dashed line boxes. Scale bar: 50 and 10 μm. B) Confocal images of WT CAL51 cells grown on a 2 kPa soft substrate coated with collagen (left) or fibronectin (right). Cells were stained with DAPI (blue), Pha-488 (green), and YAP (red). Magnified images are displayed inside gray dashed line boxes. Scale bar: 50 and 10 μm. C) 4-hour cellular uptake of PS200 (light gray) and PS900 (dark gray) in WT cells grown on a stiff polystyrene substrate coated with collagen or fibronectin. Two-way ANOVA with Sidak’s correlation for multiple comparisons. n = 3; ns, non-significant. D) 4-hour cellular uptake of PS200 (light gray) and PS900 (dark gray) in WT CAL51 grown on a 2 kPa soft substrate coated with collagen or fibronectin. Two-way ANOVA with Sidak’s correction for multiple comparisons. n = 3; ***p < 0.001. E) Schematic representation of the mechanism proposed for the differences found in nanoparticle uptake in WT CAL51 cells grown on polystyrene substrates. The cells are well-spread and attached to the surface, with high YAP nuclear localization. F) Schematic representation of the mechanism proposed for the differences found in nanoparticle uptake in WT CAL51 cells grown on soft substrates. When cells are grown on a soft substrate with a molecular-mechanical inert coating such as collagen, YAP shuttles out of the nucleus in an inactive state, and cells appear round and poorly spread. This decrease in YAP activity leads to a significant increase in nanoparticle uptake. Conversely, fibronectin coating outplays soft stiffness substrates, activates YAP and restores the mechanical properties. Nanoparticle uptake is reduced similar to what happens on stiff polystyrene.

In another set of experiments, we assessed the global endocytosis levels using fluorescent dextran (Dx-FITC) as a control for fluid-phase endocytosis and macropinocytosis levels using pHrodo-zymosan. The results obtained *via* flow cytometer demonstrate the internalization of the respective molecules and the higher uptake in YAP -/- CAL51 compared to their WT counterpart (Figure S19a-b).

When in contact with biological media, nanomaterials and nanoparticles are opsonized by several classes of proteins, which alter their surface properties and may ultimately lead to unspecific nanoparticle uptake (*57, 58*). To rule out the effect of protein corona formation on the difference in particle uptake in YAP -/- CAL51 cells, the experiment was repeated in a serum-free medium, and the obtained results remained consistent, with a marked increase in nanoparticle binding in YAP -/- CAL51 (see fig. S20a-b).

The transcriptional activity of YAP requires its shuttling to the cell nucleus, where it acts as co-activator by interacting with context-specific transcription factors (*59*). To determine the role of YAP in repressing nanoparticle uptake in TNBC cells, YAP -/- cells were transfected with a YAP hyperactive mutant (YAP-S6A). A mock vector was used as control. The shuttling of the YAP-S6A mutant protein to the nucleus is not inhibited *via* phosphorylation by the upstream Hippo pathway effectors LATS1/2 and MOB1 (Fig. 3K). The transfection was first verified by quantifying the expression of YAP protein using western blot and confocal imaging (Fig. 3L-M). The YAP-S6A- or mock-transduced CAL51 cells were then incubated with PS200 and PS900 nanoparticles. In line with the hypothesis that YAP presence in the nucleus represses nanoparticle uptake, YAP-S6A cells exhibited reduced nanoparticle uptake after 4 h of incubation compared to YAP -/- cells transfected with a mock vector (Figure 3N). Interestingly, transfecting YAP-S6A in WT cells (YAP +/+ CAL51; see fig. S21a) did not induce any significant change in nanoparticle uptake (see fig. S21b-c). This result could be explained by the fact that parental CAL51 cells already expressed a very high basal level of YAP, and the expression of a hyperactive mutant YAP-S6A did not induce any noticeable change in cell morphology (see fig. S21d), adhesion (see fig. S21e), or transcription. RT-qPCR further corroborated this hypothesis, showing no change in mRNA levels of CYR61 and CTGF, the two main transcriptional targets of YAP, in YAP +/+ CAL51 compared to WT cells (see fig. S21f).

These findings indicate that YAP activity affects cell-nanoparticle interactions and point at YAP as potential regulator of nanoparticles internalization.

### Substrate stiffness hinders nanoparticle uptake through YAP

The stiffness of the tumor stroma has been reported to impact YAP intracellular localization and transcriptional activity (*25*), which correlates with the ability of cancer cells to metastasize, leading to treatment resistance and poor prognosis (*60*). YAP intracellular shuttling and transcriptional activity are controlled by the mechanical properties of the surrounding microenvironment, with cells grown on soft substrates (E<5 kPa) exhibiting cytosolic YAP localization while the protein moves to the nucleus on stiffer substrates (E>10 kPa) (*61*). Additionally, the sensitivity of YAP to ECM components such as collagen and fibronectin has been previously documented (*9*), with fibronectin accumulation during ECM remodeling triggering YAP nuclear shuttling, independently of substrate stiffness (*18, 62*). To investigate the influence of substrate stiffness and ECM composition on nanoparticle internalization in TNBC CAL51 cells through YAP, WT -/- and YAP -/- cells were cultured on standard polystyrene plates (PSS, Young’s modulus in GPa range) or soft surfaces with a defined Young’s modulus of 2 kPa coated with either collagen or fibronectin. Confocal microscopy was used to quantify YAP subcellular localization in TNBC cells in response to substrate stiffness or ECM composition. First, we found that in the presence of collagen, the soft substrate hindered YAP nuclear localization, while the presence of fibronectin reversed this effect and restored YAP shuttling to the nucleus of cells grown on a soft substrate (Fig. 4A-B), indicating that indeed the biochemical cues from fibronectin were able to overcome the effects of the mechanical properties of the substrate under the given experimental conditions. We then investigated the relationship among substrate stiffness, YAP activity and nanoparticle uptake in TNBC cells by incubating cells cultured on soft or stiff substrates with PS200 and PS900 nanoparticles for 4 h. Our results showed that CAL51 WT cells on 2 kPa a soft substrate coated with collagen (low YAP activity) exhibited increased nanoparticle uptake, whereas this phenomenon was not observed on stiff PSS or 2 kPa substrates coated with fibronectin (high YAP activity, Fig. 4C-F). Conversely, YAP -/- CAL51 cells showed no change in nanoparticle internalization on either PSS or 2 kPa substrates coated with collagen or fibronectin (see fig. S22). These results suggest that YAP mechanical displacement from the nucleus could be an effective way to augment nanoparticle uptake in TNBC cells.

### YAP promotes ECM network deposition and cell-nanoparticle interactions in a 3D *in vitro* model of TNBC

The efficiency of endocytosis is tightly linked to the composition of the ECM that surrounds cells. We hypothesized that the increase in nanoparticle uptake measured in YAP -/- cells could be explained to a less structured ECM in these cells. We first performed an RT^2^-profiler PCR array and found that many genes related to ECM-cell adhesion were significantly upregulated in WT cells compared to YAP -/- cells (Fig. 5A; see fig. S23a-b). Some of these genes are known to be directly regulated by YAP transcriptional activity, such as CTGF (CCN2). To get a deeper insight into how YAP regulates the ECM landscape, we explored network connectivity and ontological interconnections between ECM components identified in both WT and YAP -/- cells using the STRING protein-protein interaction (PPI) database with a score threshold of 0.4 (*63*). The STRING PPI analysis yielded a highly clustered network containing 39 nodes and 336 edges and with a clustering coefficient of 0.74 for WT CAL51, indicating a significantly higher number of interactions (Fig. 5B). On the opposite, the analysis of YAP -/- CAL51 cells showed a network with fewer connections, containing 38 nodes and 165 edges, and with a lower clustering coefficient of 0.49 (see fig. S23c). As expected, the GO enrichment analysis revealed significant differences in the molecular function of the ECM network between WT and YAP -/- cells. While the former cells had high scores for annotations related to ECM and extracellular structure organization (P = 2.46×10^-12^) (Fig. 5C), the latter displayed low scores for annotations related to ECM (P = 0.0043) and extracellular structure (P = 0.0078) organization (see fig. S23d). Interestingly, the analysis also revealed that YAP depletion led to the dysregulation of several classes of ECM transcripts associated with cancer or known to promote tumor growth (see fig. S24a-b) (*64*).

**Fig. 5.**
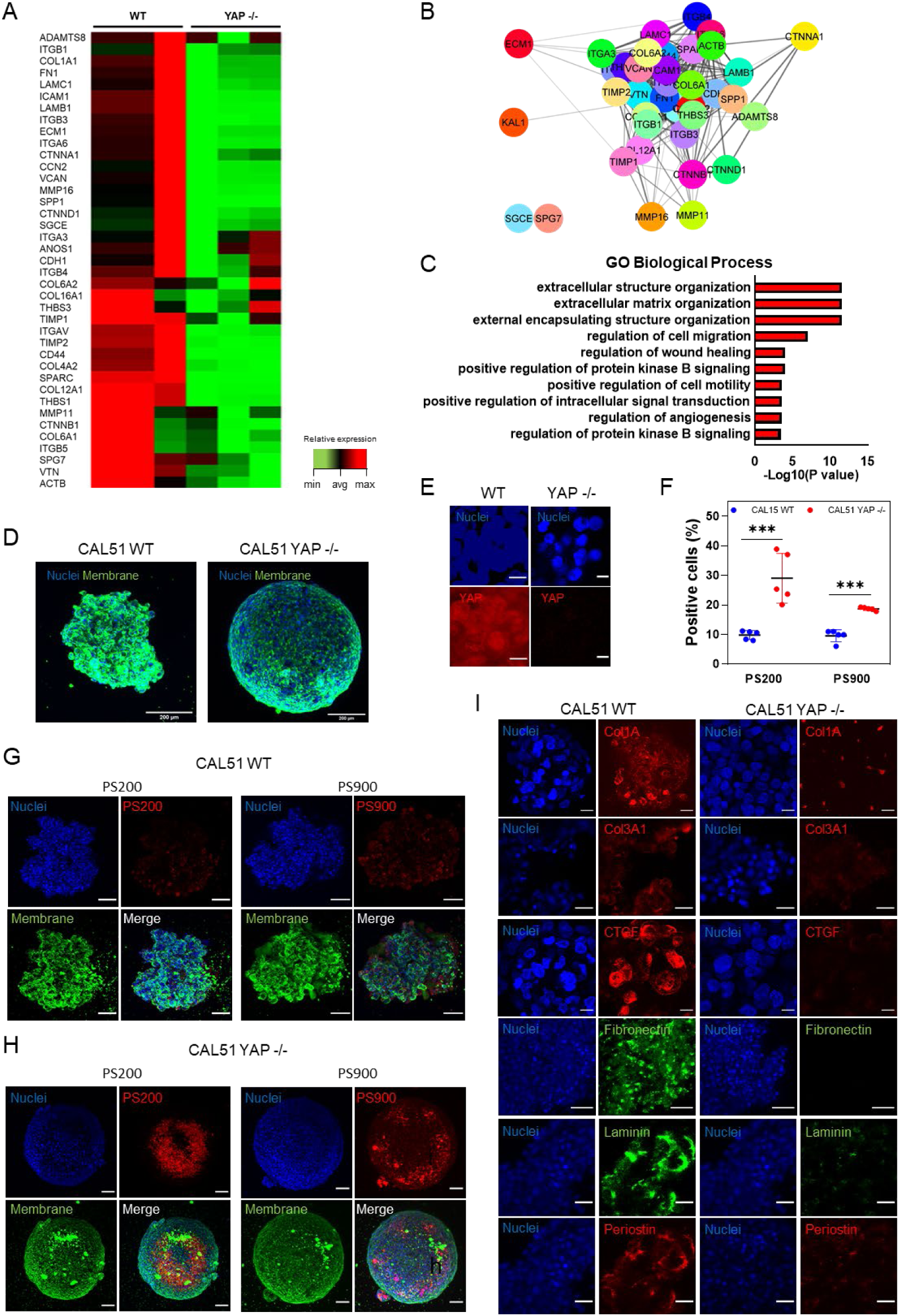
YAP knockdown affects nanoparticle binding and ECM deposition in 3D CAL51 spheroids. A) Heatmap representing the changes in extracellular matrix (ECM) and cell adhesion molecule expression in YAP -/- compared to WT CAL51 cells as obtained from RT^2^-profiler PCR array analysis (P adj < 0.05, fold change 2). B) STRING PPI network of differently expressed ECM proteins in CAL51 WT obtained from Cytoscape (P adj < 0.05, log2Fc > ǀ2ǀ, confidence cutoff 0.4). C) Bar plot representation of common enriched biological processes and pathways related to ECM network from the ENRICHR database, showing the most significantly upregulated genes in WT compared to YAP -/- CAL51 cells (P adj < 0.05, log2Fc > ǀ2ǀ). D) Z-projection images of WT (left) and YAP -/- CAL51 (right) spheroids after 5 days of culture. Cells are stained with WGA-488 (green) and DAPI (blue). Scale bar: 200 μm. E) Confocal images of the spheroids derived from WT and YAP -/- cells. Cells were stained with DAPI (blue) and YAP (red). Scale bar: 10 μm. F) 4-hour cellular uptake of PS200 and PS900 in WT (red) and YAP -/- (blue) CAL51 spheroids. Two-way ANOVA with Sidak’s correction for multiple comparisons. n = 5; ***p < 0.001. G) Confocal images of the spheroids derived from WT CAL51 cells incubated with PS200 (left) and PS900 (right) for 4 hours. H) Representative confocal images of the spheroids derived from YAP-/-CAL51 cells incubated with PS200 (left) and PS900 (right) for 4 hours. Cells were stained with WGA-488 (green) and DAPI (blue). Nanoparticles are shown in red. Scale bar: 100 μm. I) Representative confocal images of the indicated ECM components for the spheroids derived from WT (left) and YAP -/- (right) CAL51 cells. Collagen type 1 alpha (Col1A), collagen type III alpha 1 (Col3A1), connective tissue growth factor (CTGF), and periostin are stained with 2^nd^ antibody labeled with AF-555 (red); fibronectin and laminin are stained with II-antibody labeled with AF-488 (green). Scale bar: 25 μm.

Next, we aimed to validate the hypothesis that reduced ECM deposition in YAP -/- cells was responsible for increased nanoparticle uptake. We established a three-dimensional (3D) cell culture system, which resembles the complex and heterogeneous tumor microenvironment more closely than a 2D monolayer (*65*). In a 3D culture, the ECM plays a more relevant role in nanoparticle uptake than in a 2D monolayer, where ECM components are primarily deposited at the basal level. The presence of a dense network of filamentous proteins surrounding 3D spheroids has been shown to create a barrier to nanoparticle penetration and prevent physical interaction between cells and nanoparticles (*66*). We generated 3D spheroids of both WT and YAP -/- CAL51cells and investigated nanoparticle uptake using this experimental model. Briefly, the cells were seeded onto round-bottom ultra-low attachment plates and spun to promote their aggregation. As expected, after 5 days of culture, the spheroids obtained from WT or YAP -/- cells showed distinct morphologies similar to what previously described (*18*) (Fig. 5D; see fig. S25a).

Then, we investigated YAP expression in WT spheroids and found that YAP was evenly distributed in both the cytoplasm and nucleus of the cells (Fig. 5E). The live/dead assay confirmed that the cells were viable, with no detectable sign of cell death in either WT or YAP -/- spheroids (see fig. S25b). We then incubated WT and YAP -/- spheroids with PS200 and PS900 nanoparticles and quantified nanoparticle binding using flow cytometry and confocal imaging. After 4 hours of incubation, we found that YAP -/- CAL51 spheroids exhibited significantly higher nanoparticle binding than WT spheroids (Fig. 5F-H). This result was also confirmed using different nanoparticle concentrations and incubation times (see fig. S25c-d). Next, we stained WT and YAP -/- CAL51 spheroids with antibodies directed against relevant ECM proteins. The confocal microscopy analysis demonstrated that CAL51 WT cells produced a rich and multicomponent ECM composed of various proteins such as collagen, CTGF, fibronectin, laminin, and periostin, all contributing to the spheroid assembly. In contrast, YAP -/- depletion determined a stark reduction in the expression of the same ECM proteins (Fig. 5I; see fig. S26). Considering these results, targeting YAP may serve as a promising strategy for improving nanoparticle uptake into solid tumors by tuning cell membrane properties and decreasing ECM deposition.

### YAP targeting improves nanomedicine delivery to TNBC cells

After demonstrating that YAP depletion can be leveraged to increase nanoparticle uptake in TNBC CAL51 cells, we next aimed to demonstrate the therapeutic benefits of combining nanoparticle treatment with YAP targeting. To assess the efficiency of drug delivery, we chose liposomes (Fig. 6A), as they have a long history of success since Doxil (*46*), the first nano drug that reached the market, and have been recently used in nanoformulations to treat cancer and other diseases (*67-69*). We used a doxorubicin-loaded liposomal formulation (Doxo-NP) and evaluated its drug delivery efficiency in both WT and YAP -/- cells. Flow cytometry analysis showed significantly higher fluorescence intensity in YAP -/- compared to WT CAL51cells after 4-hour incubation with different concentrations of Doxo-NP (Fig. 6B; see fig. S27a). Confocal imaging confirmed a higher association of nanoparticles with YAP -/- cells (Figures 6C-D and S27b). Results from the WT and YAP -/- CAL51 co-culture and 3D spheroid experiments were also consistent (see fig. S27c and S28a-b).

**Fig. 6.**
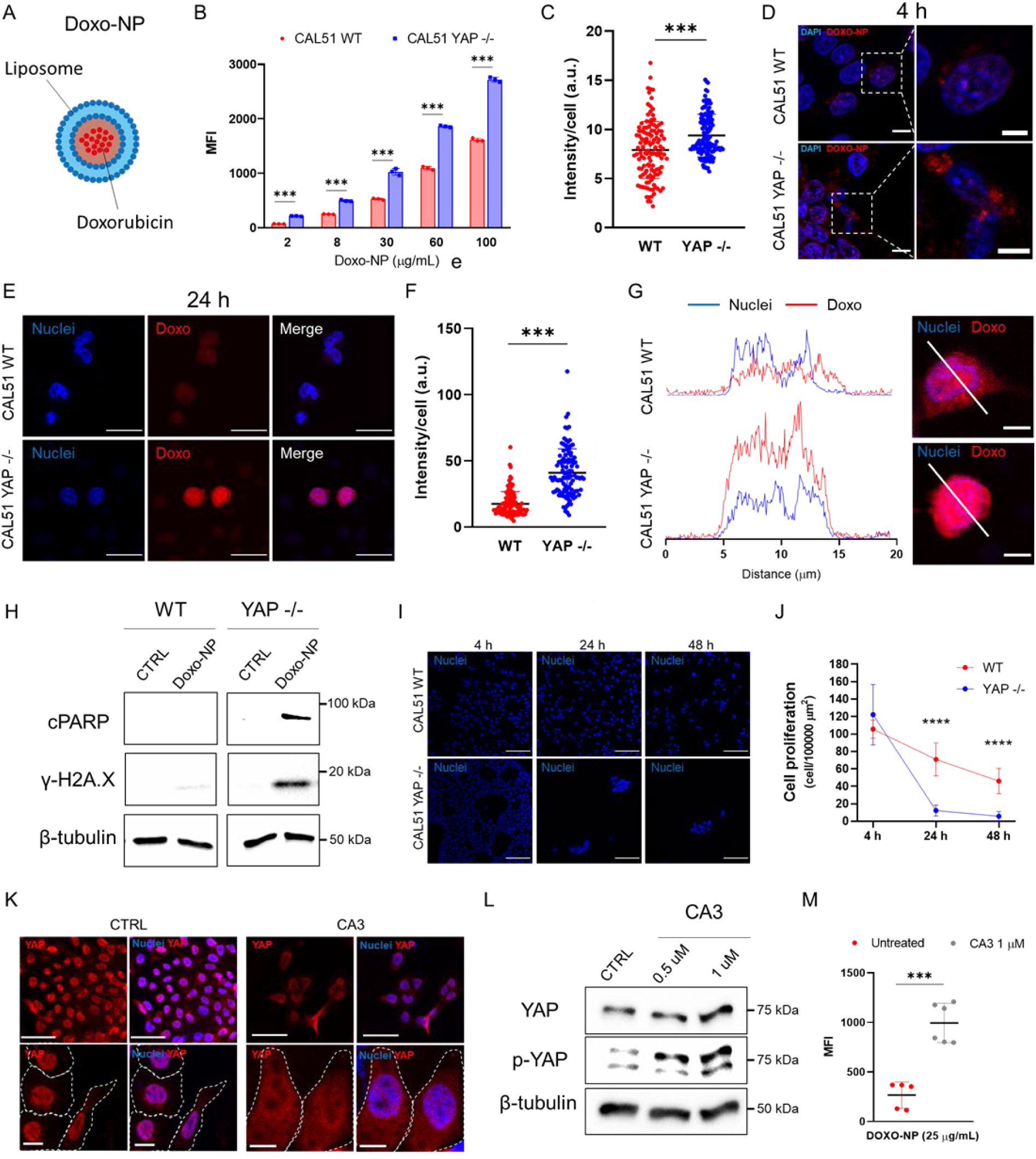
Pharmacological and genetic targeting of YAP increases the internalization of doxorubicin-loaded liposomes and improves drug delivery in TNBC CAL51 cells. A) Graphical representation of doxorubicin-loaded liposome (Doxo-NP) formulation used for drug delivery. B) Median fluorescence intensity (MFI) of Doxo-NP uptake in WT (red) and YAP -/- (blue) CAL51 as a function of nanoparticle concentration after 4-hour incubation. Two-way ANOVA with Sidak’s correction for multiple comparisons. n = 3; ***p < 0.001. C) Nanoparticle intensity per cell after a 4-hour incubation of WT and YAP -/- CAL51 cells with Doxo-NP. Unpaired t-test with Welch’s correction. n > 120; ***p < 0.001. D) Confocal images of WT (top) and YAP -/- (bottom) CAL51 cells after 4-hour incubation with Doxo-NP. Cells were stained with DAPI (blue). Nanoparticles are displayed in red. Magnified images are displayed inside white dashed line boxes. Scale bar: 10 and 5 μm. E) Representative confocal images of WT (top) and YAP -/- (bottom) CAL51 cells at 24 h after 4-hour incubation with Doxo-NP. Cells were stained with DAPI (blue). Doxorubicin is displayed in red. Scale bar: 20 μm. F) Dotplot representation of doxorubicin intensity per cell at 24 h after a 4-hour incubation of WT and YAP -/- cells with Doxo-NP. Unpaired t-test with Welch’s correction. n > 100; ***p < 0.001. G) A plot profile of doxorubicin intensity (red) co-localized with the nucleus (blue, DAPI) of WT (top) and YAP -/- (bottom) CAL51 cells. On the right, confocal images show a detailed view of the region chosen for intensity plots (white line) in WT (top) and YAP -/- (bottom) cells. Scale bar: 10 μm. H) Western blot showing the levels of cleaved PARP (cPARP) and histone H2AX (γ-H2A.X) in WT (left) and YAP -/- (right) CAL51 cells untreated (CTRL) or treated with Doxo-NP for 4 hours and collected for the analysis 48 hours post-treatment. β-tubulin was used for protein loading normalization. I) Representative confocal images of WT (top) and YAP (bottom) CAL51cells after 4 hours of incubation with Doxo-NP and 24 and 48 hours after treatment with the nanoparticles. Nuclei were stained with DAPI. Scale bar: 100 μm. J) Cell proliferation plot expressed as number of cells per surface area for WT (red line and circle) and YAP -/- (blue line and squares) CAL51cells at 0, 24, and 48 hours after 4-hour Doxo-NP treatment. Two-way ANOVA with Sidak’s correction for multiple comparisons. n = 3; ***p < 0.001. K) Representative confocal images of untreated (CTRL, left) or CA3-treated WT cells (1 μM CA3, left) for 12 hours. Cells were stained with YAP (red) and the nuclei were counterstained with DAPI (blue) and. The cell perimeter is highlighted with a dashed white line in magnified images (bottom). Scale bar: 50 and 10 μm. L) Western blot showing the levels of YAP and phospho-YAP (p-YAP) in WT CAL51 untreated (CTRL) or treated with 0.5 and 1 μM CA3 inhibitor for 12 hours. β-tubulin was used for protein loading normalization. M) MFI after a 4-hour incubation of CAL51 WT cells with Doxo-NP without treatment (red) or after treatment with 1 μM CA3 inhibitor for 12 hours (gray). Unpaired t-test with Welch’s correction. n = 2; ***p < 0.001.

Next, we extended the treatment to 24 hours and used confocal imaging to show that doxorubicin accumulated more in the nuclei of YAP -/- cells as compared to WT cells (Fig. 6E-G). Additionally, we performed western blot analysis 48 hours post-treatment with antibodies directed against cPARP and γ-H2AX and detected increased expression of both proteins in YAP -/- CAL51 cells following Doxo-NP treatment. This effect was blunted in WT cells treated with the same NPs (Fig. 6H). High toxicity was observed in YAP -/- cells at 24- and 48-hours post-treatment, with a considerably lower number of live cells per well compared to WT CAL51 (Fig. 6I-J).

Finally, we investigated if the pharmacological inhibition of YAP could be exploited to enhance Doxo-NP uptake in TNBC CAL51 cells. To this purpose, small molecule CA3 (1 μM) was used to inhibit YAP activity in breast tumor cells for 12 h (*70*) and showed no significant toxicity (see fig. S29a). Confocal images showed that CA3 treatment caused YAP to shuttle from the nucleus to the cytoplasm (Fig. 8K), which was confirmed by the western blot analysis revealing CA3-induced phosphorylation of YAP (Fig. 6L). Importantly, treatment with CA3 followed by a 4-hour incubation with Doxo-NP significantly increased nanoparticle binding and internalization (Fig. 6M; see fig. S29b). It also increased the toxicity of the treatment with the nanoformulation alone due to increased nanoparticle internalization and drug release (see fig. S29c). In light of these results, the inhibition of YAP using suitable drugs may be a promising strategy to improve cancer cell toxicity when combined with nanomedicine for the treatment of TNBC.

## Discussion

Due to its pro-tumorigenic role as an oncogene, YAP has been proposed as a target to halt cancer progression (*71*). In this study, we showed that YAP depletion in the CAL51 cell line of TNBC significantly alters the mechanical properties, surface area, and adhesion of the cells. Moreover, we demonstrated that YAP controls the genetic landscape of CAL51 cells by impacting the transcription of genes involved in membrane organization and endocytosis. RNA-seq analysis revealed differential expression of several genes involved in cell-substrate adhesion, actin-membrane linkage, ECM production, contraction, membrane tension and organization in YAP -/- relative to CAL51 WT cells. Notably, we observed that YAP depletion determines the upregulation of several genes that positively regulate endocytosis and that may contribute to the internalization of nanoparticles. In fact, from the RNA-seq data analysis (Fig. 2), we identified different transcripts associated with endocytosis that may account for the differences in the nanoparticle uptake in WT and YAP -/- cells. Interestingly, we found that CAV1, a negative regulator of endocytosis, was strongly upregulated in WT CAL51 cells (log2Fc 5.57) (see fig. S5). *CAV1* has been demonstrated to inhibit dynamin-dependent raft-mediated endocytosis, possibly acting as an endocytosis inhibitor even beyond the caveolae pathway (*72*). On the other hand, we found *DAB2*, an adapter protein that functions as a clathrin-associated sorting protein (CLASP) and that is essential for clathrin-mediated endocytosis, to be significantly upregulated in CAL51 YAP -/- cells (log2Fc 5.8) (Fig. 2D; see fig. S6). *DAB2* functions as a positive regulator of endocytosis by binding and assembling clathrin while recruiting plasma membrane components (e.g., phosphatidylinositol 4,5-bisphosphate) and receptor proteins to clathrin-coated pits (*73*). Additionally, *DLL1*, a transmembrane protein that generates mechanical force dependent on the clathrin-related endocytosis and interacting with actin filaments, was highly upregulated in CAL51 YAP -/- cells (log2Fc 7.5), contributing to endocytosis as a positive regulator (see fig. S6). Another gene identified in the RNA-seq analysis and overexpressed in YAP -/- cells included *DNM1*, which encodes Dynamin 1 and is involved in clathrin-mediated endocytosis (log2Fc 2.6), acting as a positive regulator of endocytic processes (see fig. S6).

Recent studies have shown a connection between YAP and the endocytic machinery in the cytoplasm (*74*). These interactions have long been linked to protein turnover *via* a degradation pathway alternative to the Hippo pathway, LATS1/2 phosphorylation, and proteasome. However, as our understanding of the feedback between the plasma membrane domain and mechanosensing effectors advances, the cytoplasmic pool of YAP directly interacting with proteins of the membrane, vesicles, and organelles may uncover novel and unexpected functions related to the control of organelles trafficking. Our findings suggest that YAP may play a leading role in the regulation of endocytic processes; however, its role in the homeostasis of cytoplasmic vesicles and organelles remains unclear and warrants further investigation of the molecular pathways underlying these interactions.

Throughout the study, we found that YAP knockout led to changes in cell physical and biological properties, resulting in increased nanoparticle uptake that was exclusively linked to YAP activity, not the size or surface coating of the nanoparticles. Additionally, in a co-culture system, where WT and YAP -/- CAL51 cells were seeded together, cells in which the Hippo effector had been genetically depleted showed a higher association rate with nanoparticles compared to WT cells, suggesting a possible mechanotargeting effect where cell mechanics plays a key role in bio-nano interactions. While the substrate and cell mechanics are often intertwined in bio-nano interactions and nanoparticle uptake processes (*75*), our study shows that YAP activity overrides substrate mechanics in controlling nanoparticle internalization. Indeed, irrespective of the Young’s modulus of the surface where the cells were grown, YAP activity was the key determinant in nanoparticle uptake. Specifically, an increase in YAP activity through cell-ECM interactions such as integrin-fibronectin did not enhance nanoparticle uptake even on low-stiffness substrates. Furthermore, in 3D spheroid cultures, the mechanical state of cell-cell interaction greatly depended on ECM production and deposition, which could, in turn, impact nanoparticle association and penetration within the spheroids. As a result, nanoparticles tend to associate more with spheroids derived from YAP -/- than WT cells. These data indicate that the intracellular activity of YAP and related mechanosensing proteins, not substrate mechanics, play an active role in nanoparticle internalization.

Recently, a linkage between intracellular molecular pathways and endocytosis has been established. De Belly et al. found that increased fluid-phase uptake and endocytosis are dependent on the regulation of cell membrane tension by intracellular pathways that involve the activity of β-catenin, RhoA, and ezrin-radixin-moesin (ERM) proteins (*76*). When these components are active, cells have high membrane tension, resulting in suppressed endocytosis. Degradation of β-catenin appears to reduce RhoA activity and ERM phosphorylation, lowering membrane tension. The study highlights the role of RhoA GTPase in stabilizing actin cytoskeleton anchorage to the cell membrane through YAP, thus increasing cell tension. Interestingly, our RNA-seq analysis revealed that ezrin and moesin, two ERM complex components, were significantly upregulated in CAL51 WT compared to YAP -/- cells (log2Fc 2.49 and 1.77, respectively, P adj < 0.05), revealing a close connection between the cytoskeleton, cell-substrate adhesion, and membrane tension in WT cells. Thottacherry et al. reported a decrease in membrane tension leading to increased endocytosis through the CLIC/GEEC pathway, demonstrating that this response was directly controlled by mechanical input from vinculin (*77*). Remarkably, the authors observed that in vinculin-depleted cells, fluid-phase uptake was higher than in WT cells. Our study shows that YAP depletion profoundly affects focal adhesion formation, and the subsequent rearrangement at the cell-substrate adhesion site likely reflects overall cell membrane tension and differences in the nanoparticle uptake rate between CAL51 WT and YAP -/- cells. Further studies are needed to fully elucidate the connection between YAP and the key players of endocytic pathways.

Nevertheless, our findings suggest that inhibiting YAP can help improve nanomedicine-mediated drug delivery, serving as a suitable strategy to facilitate the intracellular delivery of drugs such as doxorubicin. Indeed, the increased uptake of nanoparticles into YAP -/- cells resulted in a higher accumulation of the drug in the nucleus, as a direct result of YAP inhibition.

Remarkably, a recent study has shown that cancer cell migration and extravasation, critical for forming distant metastasis by entering the vasculature, depends on the cell sensitivity to shear stress and the activation of Rho GTPases such as RhoA (*78*). In particular, the inhibition of Rho-associated protein kinase (ROCK) was found to abolish cell sensitivity to shear stress and promote their extravasation. As ROCK function correlates with the activity of YAP and ERM proteins as well as membrane tension, our findings suggest that migrating cells may be more susceptible to nanoparticle targeting due to changes in membrane organization and forces. This evidence suggests that leveraging mechanosensitive pathways in cancer cells may be a potential strategy to selectively target highly migrating cells and cancers with pro-metastatic features.

Cell mechanobiology contributes to cell-nanoparticle interactions, and the functions of its principal pathways and effectors may improve nanoparticle delivery to cancer cells. The mechanobiology pathways identified here may be leveraged to ameliorate the design of nanoparticles, focusing on the development and characterization of nanomaterials that are able to interact with the cell membrane in more efficient ways. In this study, we have demonstrated that YAP co-transcriptional activity hinders nanoparticle binding and internalization. As we explored along the paper, several reasons may account for this association and can be ascribed to the role of YAP in: i) directing the transcription of genes involved in cell adhesion and mechanosensing; ii) affecting the genetic landscape of endocytic pathways by transcriptionally suppressing proteins involved in endocytosis; iii) perturbing cell membrane tension and organization, thus promoting its deformation and facilitating the formation of endocytic vesicles; iv) producing an abundant and dense ECM network that may ultimately hamper nanoparticle diffusion (see fig. S30).

Nonetheless, further research is needed to fully understand the relationship between cellular mechanosensing and nanomaterials. Establishing whether YAP is directly involved in the endocytic process remains - in fact – difficult, due to the substantial overlap between cell mechanics and function. Moreover, while the role of YAP in cell nucleus has been extensively characterized, the activity of its cytoplasmic pool is still poorly understood and further investigations are now warranted in this direction. Finally, notwithstanding YAP depletion affects well characterized key features of the tumor microenvironment that contribute to limit the application of nanoparticles in cancer treatment, the translational potential and clinical relevance of this approach remain to be confirmed.

In conclusion, by genetically, mechanically or pharmacologically targeting YAP, we show that it is possible to increase nanoparticle association and internalization in TNBC cells, highlighting the role of mechanobiology in shaping the fate of bio-nano interactions in cancer cells. We demonstrate that blocking YAP activity may be used to increase the delivery of nano drugs, paving the way for novel combinatorial therapies suited to tackle cancer tumorigenicity while simultaneously enhancing the delivery of anti-cancer nanotherapeutics. This work opens up new avenues for selectively tuning cell-nanoparticle interactions by targeting molecular processes that differentiate between cancer cells and their healthy counterpart, thus improving both the delivery and specificity of nanotherapy. To assess the pre-clinical and clinical value of such nanotherapies, we propose an alternative fundamental mechanism for nanoparticle entry into the cells. A deeper understanding of the cell mechanobiology pathways in bio-nano interactions and the search for new targets and drugs to modulate their functions could accelerate the development of advanced next-generation nanotherapies that would address the challenges posed by nanomedicine concerning targeted and selective drug delivery.

## Materials and Methods

### Materials

Polystyrene carboxylated nanoparticles 0.9 and 0.204 μm (W090CA and 83000520100290, respectively, Thermo fisher); µ-Dish 35 mm, 1.5 ESS (81291, Ibidi); CytoSoft Imaging, 24-well Plate, Elastic Modulus 2 kPa (5185-1EA, CellSystem); bovine plasma fibronectin (F1141, Merck); Collagen I, rat tail (A1048301, Thermo Fisher); Dulbecco’s Modified Eagle Medium (DMEM, Merck); Opti-Mem (31985062, Thermo Fisher); 6-well plate (30006, SPL Life Sciences); 12-well plate (30012, SPL Life Sciences), 24-well plate (30024, SPL Life Sciences); 96-well ultralow attachment plates (7007, Corning); µ-slide 8-well glass bottom dish (80807, Ibidi), Tissue culture dish 40 mm (93040, Ibiotech); Alexa Fluor™ 647 Phalloidin (A22287, Thermo Fisher); Alexa Fluor™ 488 Phalloidin (A12379, Thermo Fisher); Wheat Germ Agglutinin, Alexa Fluor™ 647 Conjugate (W32466, Thermo Fisher); Wheat Germ Agglutinin, Alexa Fluor™ 488 Conjugate (W11261, Thermo Fisher); pLX304 (Addgene plasmid # 25890; http://n2t.net/addgene:25890; RRID:Addgene_25890); YAP1 (S6A) - V5 in pLX304 (Addgene plasmid # 42562; http://n2t.net/ addgene: 42562; RRID:Addgene_42562); FuGENE HD (E2311, Promega); Lipofectamine 3000 (L3000001, Thermo Fisher); 10% Mini-Protean TGX Precast Protein gel (4561033, Bio-Rad); protease and phosphatase inhibitor cocktails (PPC1010, Merck); RIPA buffer (R0278, Merck); HRP-conjugated anti-rabbit and HRP-conjugated anti-mouse (RABHRP1 and RABHRP2, Merck); MOWIOL 4-88 Reagent (475904, Merck); LysoTracker Deep Red (L12492, Thermo Fisher); High Pure RNA Isolation Kit (11828665001, Roche); RT² Profiler™ PCR Array Human Extracellular Matrix & Adhesion Molecules (Ref. PAHS-013Z); LIVE/DEAD Viability/Cytotoxicity Kit (L3224, Thermo Fisher); CellTrace CFSE Cell Proliferation Kit (C34554, Thermo Fisher); Dextran, Fluorescein, 10,000 MW (D1821, Thermo Fisher); pHrodo Green Zymosan Bioparticles (P35365, Thermo Fisher); Opti-Link Carboxylate-Modified Particles (83000520100290 and W090CA, Thermo Fisher); TAMRA cadaverine (92001, Biotium); Fluorescein Cadaverine (A10466, Thermo Fisher); Flot-A-Lyzer G2 dialysis device (300 kDa cutoff) (G235036, Merck); CellTracker Blue CMAC Dye (C2110, Thermo Fisher); N-Hydroxysuccinimide (804518, Merck); Chlorpromazine hydrochloride (sc-202537, Santa Cruz); Nystatin (sc-2122431, Santa Cruz); Cytochalasin D (C8273, Merck); CA3 (CIL56) (S8661, Selleckchem); EDC-hydrochloride (59002, VWR); sodium cacodylate trihydrate (20840, Merck); glutaraldehyde 25% (354400, Merck); 4′,6-Diamidine-2′-phenylindole dihydrochloride (10236276001, Merck); Dox-NP (300112, Avanti Polar Lipids); tannic acid (403040, Merck); Iron chloride tetrahydrate (380024, Merck); mouse anti-YAP (4912, Cell Signaling); mouse anti-Vinculin (V9131, Merck); anti-Col1A1 (91144S, Cell Signaling); anti-Col3A1 (PA5-27828, Thermo Fisher); anti-CTGF (Ab5097, Abcam), anti-Fibronectin (F3648, Merck); anti-Laminin (SAB4200719, Merck); anti-Periostin (49480, Santa Cruz); rabbit anti-YAP (14074, Cell Signaling); mouse anti-β-tubulin (T8328, Merck); mouse anti-vinculin (V9131, Merck); rabbit anti-pYAP S127 (D9W21, 13008, Cell Signaling); rabbit anti-pH2A.X (2577, Cell Signaling); rabbit anti-cPARP (5625S, Cell Signaling). All antibodies used for western blotting were diluted at a ratio of 1:100. Antibodies for immunohistochemistry were used according to the specifications in Table 1 in the *Supporting information*.

### Polystyrene nanoparticle fluorescent labeling

Polystyrene carboxylated nanoparticles with a size of 0.9 and 0.204 μm were labeled with TAMRA cadaverine or Fluorescein cadaverine. Briefly, the particles were resuspended in 5 mL MES buffer (50 mM, pH=6.04) at a final concentration of 20 mg/mL and incubated with 52 μM/mL EDC-hydrochloride and 5.2 μM/mL N-hydroxysuccinimide for 1 hour in an ice bath under magnetic stirring. Afterward, TAMRA cadaverine or fluorescein cadaverine was added, and the mixture was brought to a final concentration of 50 μM and left to react overnight (O.N.) at room temperature (RT). The day after, the particles were collected, centrifuged (5,000 rpm for 10 min for 0.9 μm particles, 12,000 rpm for 15 min for 0.204 μm particles), and washed five times with distilled water. Subsequently, the particles were dialyzed against distilled water for 72 hours using a Flot-A-Lyzer dialysis device (300 kDa cutoff).

### Tannic Acid (TA)/Fe^III^ Coating on Nanoparticles

The MPN coating process was performed after fluorescein cadaverine functionalization of the nanoparticles; 250 μL of the particles were dispersed in an equal amount of distilled water, and 15 μL (PS 0.9 μm) or 5 μL (PS 0.204 μm) of FeCl3·6H2O (37 mM) added under sonication. After 1 min, 15 μL (PS 0.9 μm) or 5 μl (PS 0.204 μm) of tannic acid (TA) solution (24 mM) solution was added, and the mixture was left to react for 5 min under sonication. Afterward, the pH was raised by adding 500 μL of MOPS (20 mM, pH 7.4), and the suspension was vortexed for 5 min to ensure adequate MPN film formation and adherence. The particles were then centrifuged (5,000 rpm for 10 min for 0.9 μm particles, 12,000 rpm for 15 min for 0.204 μm particles) and washed five times with distilled water. Finally, the particles were resuspended in 250 μL of distilled water.

### Generation of YAP mutant CAL51 lines

The YAP -/- CAL51 lines were generated using CRISPR/Cas9 technology as described previously.(*18*) Briefly, the process involved designing a guiding RNA to target exon 1 of the YAP1 gene, which is common in all nine YAP1 splicing variants. Two sets of complementary single-stranded DNA oligonucleotides (YAP1_R1: 50-CACCgtgcacgatctgatgcc-30, YAP1_R2: 50-AAACccgggcatcagatcgtgcac-30, YAP1_F1: 50-CACCGcatcagatcgtgcacgt-30, YAP1_F2 50-AAACcggacgtgcacgatctgatgC-30) were then cloned into pSpCas9(BB)-2A-GFP (PX458) and transfected into Cal51 cells using the FuGENE HD transfection reagent according to the manufacturer’s protocol. The next day, green fluorescent protein (GFP)-positive cells were FACS-sorted (MoFlo Astrios, Beckman Coulter, California, USA) as single cells into a 96-well plate and clonally propagated. Genomic DNA was sequenced from both sides to map the size of the deletion (sequencing primers: 50-gattggacccatcgtttgcg-30, 50-gtcaagggagttggagggaaa-30, 50-gaagaaggagtcgggcagctt-30, 50-gagtggacgactccagttcc-30).

### Cell Culture

Wild-type (WT) CAL51 (gift from Dr. L. Krejčí, Department of Biology, Masaryk University) and mutant cell line were cultured in Dulbecco’s Modified Eagle Medium (DMEM) containing 10% Inactivated Fetal Bovine Serum (FBS), 1% Penicillin-Streptomycin (PS), and 1% Glutamine at 37 °C, 95% humidity, and 5% CO2. The cells were split every 2-3 days before reaching confluence, and only the CAL51 cells with a passage number of < 20 were used for all the experiments. Spheroids were generated by seeding 3,000 cells per well in a volume of 100 μL in 96-well ultralow attachment plates and aggregating them *via* centrifugation at 1000 rpm for 5 min. The spheroids were left to grow for 5 days with media replacement every 2 days. The cell proliferation assay was performed using the CellTrace CFSE Cell Proliferation Kit (C34554, Thermo Fisher). Cells were labeled with the dye according to the supplier’s protocol and seeded onto a 24-well plate (30024, SPL Life Sciences) for 24 hours. Flow cytometry analysis was performed using unstained cells as a control.

### Atomic force microscopy (AFM) measurements

Cells were seeded onto a 40 mm tissue culture (93040, Ibiotech) at a concentration of 100,000 cells per dish for 24 h. Force maps were measured using a bio-AFM Nanowizard 4XP (Bruker-JPK, Germany) placed on a Leica DMi 8 inverted microscope with a 10x objective (Leica, Germany). A 5.73 µm melamine sphere (microParticles, Berlin, Germany) was attached to a soft tipless cantilever SD-qp-CONT-TL (NanoWorld, Neuchâtel, Switzerland) using epoxy resin. A plastic Petri dish (TPP, Trasadingen, Switzerland) containing either distilled water for calibration or cell culture was placed on a motorized stage with a Petri dish heater pre-heated to 37 °C. Before each experiment, the laser reflection sum was maximized, and the laser detector was centered. Immediately after, probe sensitivity and stiffness were determined using the thermal noise method in Bruker-JPK software. AFM settings were kept constant for each sample, with a setpoint in the range of 0.2-0.8 nN relative to the baseline to always maintain indentation depth up to 2.8 µm, Z-length at 10 µm, speed of recording at 20 µm s-1, and sample rate at 5 kHz. Each force map consisted of 64×64 or 32×32 FDCs (Force-distance curves) covering an area of single or multiple cells, and Younǵs modulus was calculated from the FDCs by fitting the DMT model (*79*) in AtomicJ software (*80*).

### Correlative Probe and Electron Microscopy measurements by LiteScope^TM^

Correlative Probe and Electron Microscopy (CPEM) images were acquired using an AFM LiteScope (NenoVision, Czech Republic) integrated with a Versa 3D Dual Beam SEM (Thermo Fisher Scientific, Czech Republic). This setup allowed for the identification of particles on/in cells and navigation of the tip to areas of interest by electron imaging. SEM and AFM topography images were simultaneously acquired in NenoView software, showing differences in material and topological features. Topography was measured using a frequency-modulated tapping regime with an Akiyama probe (Nanosensors), a self-sensing probe with a visible apex. At the same time, the electron beam was focused near the AFM tip to acquire secondary electron signals from the Ion Conversion and Electron (ICE) detector. All images were acquired with 512 points and a scanning speed between 15 µm/s and 2 µm/s, depending if large or small scanning fields were acquired, with a primary electron acceleration voltage of 10kV and current of 1.5pA. The offset between the SE image and AFM topography was then removed in NenoView software, using the same pixel size for correlation. Final post-processing was performed in Gwyddion software (Czech Republic).

### Scanning electron microscopy

WT and YAP -/- CAL51 cells were cultured on coverslips for 2 days, fixed with a solution of 3% Glutaraldehyde in 100mM Cacodylate buffer, and dehydrated in a series of increasing ethanol concentrations of 30%, 50%, 70%, 80%, 96%, and 100%. The samples were mounted on aluminum stubs, sputter-coated with Palladium (JEOL JFC-1300, Tokyo, Japan), and visualized using a Benchtop Scanning Electron Microscope JEOL JCM-6000 (Tokyo, Japan).

### Transmission electron microscopy

WT or YAP -/- CAL51 cells were grown on 24-well plates and left for 12 h to allow for adhesion, then incubated with nanoparticles for 4 h in a growth medium. Afterward, the cells were washed, detached, and seeded for an additional 4 h. Fixation was done by 1.5% PFA + 1.5% glutaraldehyde diluted in DMEM high glucose culture media for 1 h, then washed with 0.1 M cacodylate buffer (pH 7.4) for 3 times 5 min each, and incubated overnight at 4 °C in cacodylate 0.1 M with 1.5% glutaraldehyde. The next day, samples were rewashed 3 times with cacodylate buffer, and incubated at 4 °C in cacodylate buffer until sending the samples, protected from light. The obtained cell pellet was post-fixed for 1.5 h with 1% osmium tetroxide in 0.1 M cacodylate buffer and washed for 3 times with 0.1 M cacodylate buffer, 10 min each, then stained with 1% uranyl acetate in milli-Q water overnight at 4 °C, followed by washing in milli-Q water. The samples were then dehydrated in an ascending EtOH series using solutions of 70%, 90%, 96% and 3 times 100% for 10 min each, incubated in propylene oxide (PO) 3 times for 20 min before incubation in a mixture of PO and Epon resin overnight, and incubated in pure Epon for 2 h and embedded by polymerizing Epon at 68 °C for 48 h. Ultra-thin sections of 70 nm were cut using a Leica Ultracut EM UC 6 Cryo-ultramicrotome. TEM images were collected with a JEOL JEM 1011 electron microscope and recorded with a 2 Mp charge coupled device camera (Gatan Orius).

### Isolation of RNA and PCR analysis

Total RNA was isolated using High Pure RNA Isolation Kit (Roche, Switzerland), according to the manufacturer’s protocol. Complementary DNA was synthesized using the RT^2^ First Strand Kit (SABiosciences, Frederick, USA). The expression levels of genes involved in ECM and cell adhesion were analyzed using RT^2^ Profiler PCR Arrays (Qiagen). Reverse transcription polymerase chain reaction (RT–PCR) was performed on the LightCycler 480 Real-Time PCR System (Roche, Basel, Switzerland), and the cycling parameters were 1 cycle at 95 °C for 10 min, followed by 45 cycles at 95 °C for 15 s and 60 °C for 1 min. The normalization of gene expression levels was performed using an internal panel of housekeeping genes provided by the manufacturer. The PCR-array data were analyzed online using the tools available on the manufacturer’s website (http://www.sabiosciences.com/pcrarraydataanalysis.php). Genes with a high coefficient of variation among replicas or very low expression (35 < Ct > 40) were excluded from the analysis. The results are presented as heatmaps of quantification cycles (Ct) and graphs of mean ± standard deviation (s.d.) values of fold regulation, based on three samples per experimental condition.

### RNA Sequencing (RNA-seq) and Data Analysis

The library was prepared using the NEBNext® Ultra™ II Directional RNA Library Prep Kit for Illumina® with NEBNext® Poly(A) mRNA Magnetic Isolation Module and NEBNext® Multiplex Oligos for Illumina® (Dual Index Primers Set 1). The kits were employed according to the manufacturer’s protocol, and 200-300 ng of total RNA was used as input to prepare the library. Sequencing was performed on an Illumina NextSeq 500 using the NextSeq 500/550 High Output v2 kit (75 cycles). Single-end 75bp sequencing was carried out in multiple sequencing runs until all samples had at least 30 million passing filter reads. Fastq files were generated using bcl2fastq software without any trimming. The quality of the raw sequencing data was assessed using FastQC (https://www.bioinformatics.babraham.ac.uk/projects/fastqc/) and aligned to the hg38 reference genome using TopHat2 aligner. Raw gene counts were calculated from reads mapping to exons, summarized by genes using the Ensembl 90 reference gene annotation (Homo sapiens GRCh38.p10, GTF) by HTSeq. Differential gene expression was determined using the DESeq2 Bioconductor package. Genes were considered differentially expressed if a Benjamini-Hochberg adjusted P-value was ≤ 0.05 and log2 fold-change (log2FC) was ≥ 1.5. Biological term classification and gene cluster enrichment analysis were performed using the clusterProfiler package. All computations were performed using BioJupies (*81*). Enrichment analysis and ranking were performed using Enrichr, and the most significant annotations were downloaded from the available repository (https://maayanlab.cloud/Enrichr/) (*82-84*). The Gene Ontology (GO) categories were downloaded from the AmiGO 2 repository (https://amigo.geneontology.org).

### Cell transfection

Cells were transfected using Lipofectamine 3000. The plasmids YAP1 (S6A)-V5 in pLX304 and pLX304 were obtained from Addgene as gifts from William Hahn (plasmid 42562) and David Root (plasmid 25890), respectively (*85*). CAL51 cells were seeded onto a 6-well plate and transfected 24 h later with a preincubated mixture containing 250μL of Opti-MEM, 2.5 ng of DNA, 7.5 μl of Lipofectamine 3000, and 5 μL of P3000 reagent, added dropwise into each well. After 12 h, fresh medium was added, and cells were allowed to grow for another 8 h. The cells were then detached, seeded in a 24-well plate at a density of 200,000 cells/well, and cultured for an additional 12 h prior to incubation with nanoparticles.

### Polystyrene nanoparticle internalization studies

The day before the experiments, 200,000 cells were seeded in 500 μL of medium onto a 24-well plate. For co-culture experiments, CAL51 YAP -/- cells were labeled with CellTracker Blue CMAC Dye according to the manufacturer’s protocol. Then, 100,000 CAL51 WT and 100,000 pre-labeled CAL51 YAP -/- cells were seeded together onto a 24-well plate. For the experiments carried out using substrates with different stiffness, either a µ-Dish 35 mm, 1.5 ESS or a CytoSoft Imaging 24-well Plate with an Elastic Modulus of 2 kPa that were pre-coated with 0.1 mg/mL of collagen I rat tail or 30 μg/mL of bovine plasma fibronectin have been used. After 18 h, the cells were incubated with nanoparticles diluted at the desired concentration in the supplemented medium for 4 h (or according to the time indicated for each experiment), with 500 μL of nanoparticle dispersion added per well. The samples were then processed for downstream flow cytometry and confocal analysis according to the following protocols: for cytofluorimetric analysis, CAL51 WT or YAP -/- cells were cultured on 24-well plates and left for 12 h to adhere, then incubated with PS200 and PS900 for 4 h. The medium was removed, 200 μL of TrypLE Express Enzyme was added to each well for 5 min, and the cells were detached and washed three times until no particle signal was detected in the supernatant. The samples were analyzed using the FACSAria II (Becton-Dickinson, USA), and plots were prepared with FlowJo software V10 (Tree Star, USA). For confocal laser scanning microscopy (CSLM) analysis, after the incubation with nanoparticles, the cells were detached, washed three times with PBS, and left to adhere to a µ-slide 8-well glass bottom dish. After 4 h, the cells were processed according to the *immunohistochemistry* protocol described below.

### Mechanism of nanoparticle endocytosis study

To investigate the mechanism of nanoparticle endocytosis, the cells were cultures in a medium containing endocytosis inhibitors chlorpromazine hydrochloride (10 μg/ml), nystatin (100 U/ml), or cytochalasin D (1 μg/ml) for 2 h, then incubated with polystyrene nanoparticles for 4 h. After incubation, the cells were processed for flow cytometry. Endo-/lysosomal trafficking was also examined by adding LysoTracker Deep Red to the culture media until a final concentration of 100 nM, with the cells incubated for endo-/lysosome staining for 1 h according to the manufacturer’s protocol. The cells were then processed for immunohistochemistry analysis according to the previously described protocol.

### Doxorubicin-loaded liposome (Doxo-NP) internalization study

To evaluate the role of YAP in nanomedicine delivery studies, CA51 WT and YAP -/- cells were incubated with Doxo-NP for 4 h before examining cell-nanoparticle interactions. To evaluate the pharmacological inhibition of YAP, Doxo-NP internalization was examined in CAL51 WT cells pre-treated with 1 μM of CA3 inhibitor for 12 h. After replacing the media, the cells were incubated with Doxo-NP for another 4 h, and then analyzed by flow cytometry or CLSM, as previously described.

### Immunohistochemistry and image analysis

For immunohistochemistry, 40,000 cells per well were seeded onto a µ-slide 8-well glass bottom dish for 24 h. After each relevant experiment, the medium was removed, and the cells were washed with PBS. Before staining, the cells were fixed with 200 μL 4% paraformaldehyde in PBS for 15 min at RT, permeabilized with 0.1% Triton X-100 for 5 min, and then blocked with BSA 2.5% in PBS for 30 min. The primary antibodies were added in 200 μL PBS-BSA 2.5% solution and incubated for 2 h at RT or O.N. at 4 °C. Afterward, the secondary Alexa fluorochrome-conjugated antibodies were added, and the cells were incubated in PBS. F-actin was stained with Alexa Fluor 488 or 647-conjugated phalloidin, the membrane was stained with wheat germ agglutinin (WGA) 488- or 647-conjugated, and the nuclei were counterstained with 4’,6’-diamidino-2-phenylindole (DAPI) or Hoechst. The samples were embedded in Mowiol reagent (Merck) and visualized with a Zeiss LSM 780 or Leica TCS SP8 X confocal microscope. Z-stacks were acquired with an optimal interval suggested by the software, and a maximum intensity algorithm was applied. Images were analyzed using ImageJ (http://rsb.info.nih.gov/ij/). The primary antibodies used were mouse anti-YAP (4912, Cell Signaling), mouse anti-Vinculin (V9131, Merck), anti-Col1A1 (91144S, Cell Signaling), anti-Col3A1 (PA5-27828, Thermo Fisher), anti-CTGF (Ab5097, Abcam), anti-Fibronectin (F3648, Merck), anti-Laminin (SAB4200719, Merck), and anti-Periostin (49480, Santa Cruz), as referred in the Materials sections and in Supplementary Table 1.

### Western blotting

Cells were lysed with RIPA buffer (Merck Millipore) containing 1% protease and phosphatase inhibitor cocktails on ice, and then centrifuged at 14.000 g for 15 min at 4 °C. The supernatants were stored at 80 °C. Protein concentrations were determined using the Bicinchoninic Acid (BCA) protein assay, and 20 μg of protein per sample were loaded onto 10% polyacrylamide gels and run at 100V. Proteins were transferred to a polyvinylidene difluoride membrane using the Trans-Blot Turbo transfer system (Bio-Rad). The membranes were blocked with 5% BSA in TBST and incubated with primary antibodies diluted in 5% BSA in TBST O.N. at 4 °C. They were then probed with proper HRP-linked secondary antibodies for 1 h at RT. Chemiluminescence was detected using the ChemiDoc imaging system (Bio-Rad), and band intensities were quantified using Bio-Rad Image Lab software. The primary antibodies used were rabbit anti-YAP, mouse anti-β-tubulin, mouse anti-vinculin, rabbit anti-pYAP S127, rabbit anti-pH2A.X, and rabbit anti-cPARP.

### Minimum Information Reporting in Bio−Nano Experimental Literature (MIRIBEL)

The studies conducted here, including material characterization, biological characterization, and experimental details, conform to the MIRIBEL reporting standard for bio−nano research (*86*). A companion checklist for the MIRIBEL components is provided in the *Supporting Information* (Tables 2-4).

### Statistical analysis

Results are based on at least three replicates, and the data are presented as the mean ± s.d. The calculations were performed using GraphPad Prism v. 6.0 (San Diego, USA). For single-cell analysis, a minimum of 100 cells per sample were considered. Appropriate statistical tests were applied to the data as indicated in the figure captions for each experiment. A P < 0.05 was considered statistically significant.

## Supporting information

Cassani et al. 2023_SI

## Acknowledgments

We are grateful to František Foret for granting the access to its facility at the Department of Bioanalytical Instrumentation of the institute of analytical chemistry of Brno. We thank Jana Bartoňová, Stefania Pagliari and Vladimír Vinarský for scientific advice and technical assistance. We thank Roberto Marotta and Federico Catalano of the Electron Microscopy Facility, Fondazione Istituto Italiano Di Tecnologia for technical assistance. We acknowledge the Department of Chemical Engineering of the University of Melbourne for their support in performing part of the experiments presented in this paper. We acknowledge the CF Genomics of CEITEC supported by the NCMG research infrastructure (LM2018132 funded by MEYS CR) Bioinformatics for their support with obtaining scientific data presented in this paper. We also acknowledge CIISB, Instruct-CZ Centre of Instruct-ERIC EU consortium, funded by MEYS CR infrastructure project LM2023042 and European Regional Development Fund-Project “UP CIISB” (No. CZ.02.1.01/0.0/0.0/18_046/0015974), is gratefully acknowledged for the financial support of the measurements at the CF Nanobiotechnology. We also thank Romana Vlčková, Hana Dulová and Jana Vašíčková for their support on continuation of the study. The authors thank Ilya Demchenko from Insight Editing London for editorial support of the manuscript.

## Funding

Marco Cassani, an iCARE-2 fellow, has received funding from Fondazione per la Ricerca sul Cancro (AIRC) and the European Union’s Horizon 2020 research and innovation programme under the Marie Skłodowska-Curie Grant Agreement No. 800924. Giancarlo Forte was supported by the European Regional Development Fund— Project ENOCH (No. CZ.02.1.01/0.0/0.0/16_019/0000868). Jorge Oliver-De La Cruz and Soraia Fernandes were supported by the European Social Fund and European Regional Development Fund-Project MAGNET (CZ.02.1.01/0.0/0.0/15_003/0000492). This work was supported by Marie Curie H2020-MSCA-IF-2020 MSCA-IF-GF “MecHA-Nano”, Grant Agreement No 101031744.

## Author contributions

Conceptualization: MC, GF. Methodology: MC, SF, JODLC, JV, VH, MP, SK, JP, DD. Investigation: MC, SF. Visualization: MC. Supervision: MC, GF. Writing—original draft: MC, GF. Writing—review & editing: MC, SF, FC, GF.

## Conflict of interest

The author declares that there is no conflict of interest in connection with this article.

## Data and materials availability

All data needed to evaluate the conclusions in the paper are present in the paper and/or the Supplementary Materials. Additional data related to this paper may be requested from the authors.

## Notes

### Competing Interest Statement

The authors have declared no competing interest.

