## Supplementary material for "YAP signaling regulates the cellular uptake and therapeutic effect of nanoparticles": Cassani et al. 2023_SI

[1]International Clinical Research Center, St. Anne's University Hospital, Brno, Czech Republic. [2]NenoVision, Purkynova 649/127, Brno, 61200, Czech Republic. [3]Faculty of Mechanical Engineering, Brno University of Technology, Technicka 2896/2, Brno, 61669, Czech Republic. [4]Department of Bioanalytical Instrumentation, CEITEC Masaryk University, Brno, Czech Republic. [5]Electron Microscopy Facility, Fondazione Istituto Italiano Di Tecnologia, Via Morego 30, 16163, Genoa, Italy. [6]School of Science, RMIT University, Melbourne, Victoria, Australia. [7]Dipartimento di Scienze e Tecnologie Chimiche, Università di Roma "Tor Vergata", Via Della Ricerca Scientifica, Rome, Italy. [8]Department of Chemical Engineering, The University of Melbourne, Parkville, Victoria, Australia. [9]School of Cardiovascular and Metabolic Medicine & Sciences, King's College London, London WC2R 2LS, UK.

*Present address: Institute for Bioengineering of Catalonia (IBEC), The Barcelona Institute for Science and Technology (BIST), Barcelona, Spain.

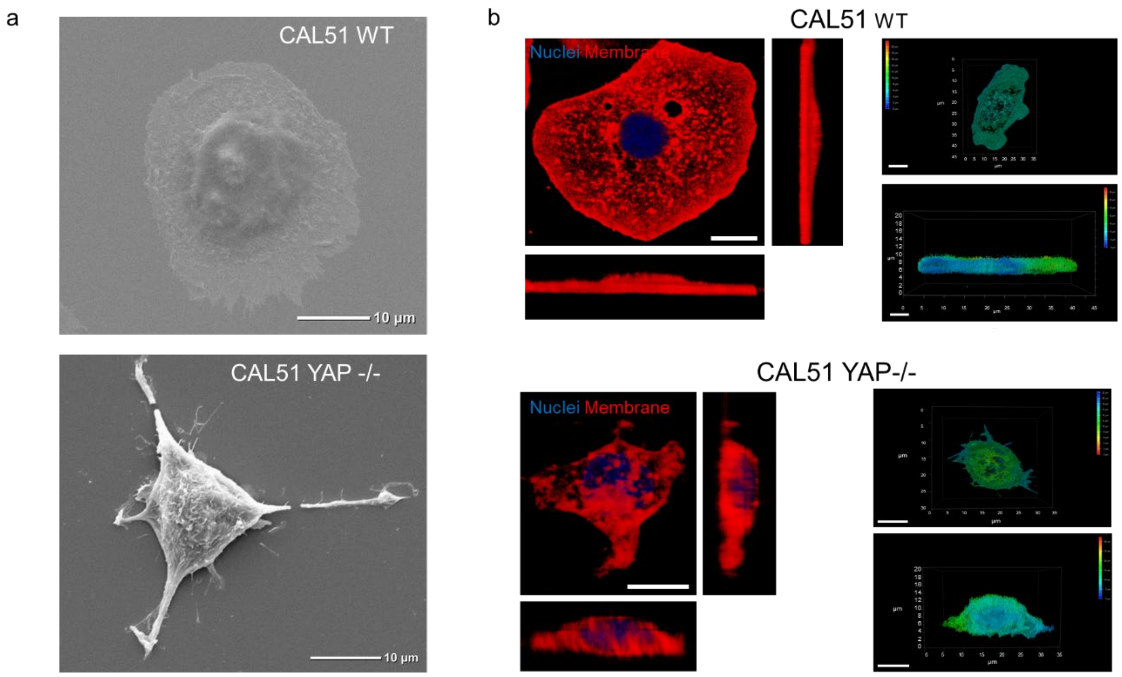

**Fig. S1.** a) Representative SEM images of WT and YAP -/- CAL51 cells. Scale bar: 10 µm. b) 3D reconstruction of WT (top) and YAP -/- (bottom) CAL51 cells. The top and lateral views are presented. The cells on the left are stained with DAPI (blue) and WGA-647 (red). The cells on the right are presented in depth color code. Scale bar: 20 µm.

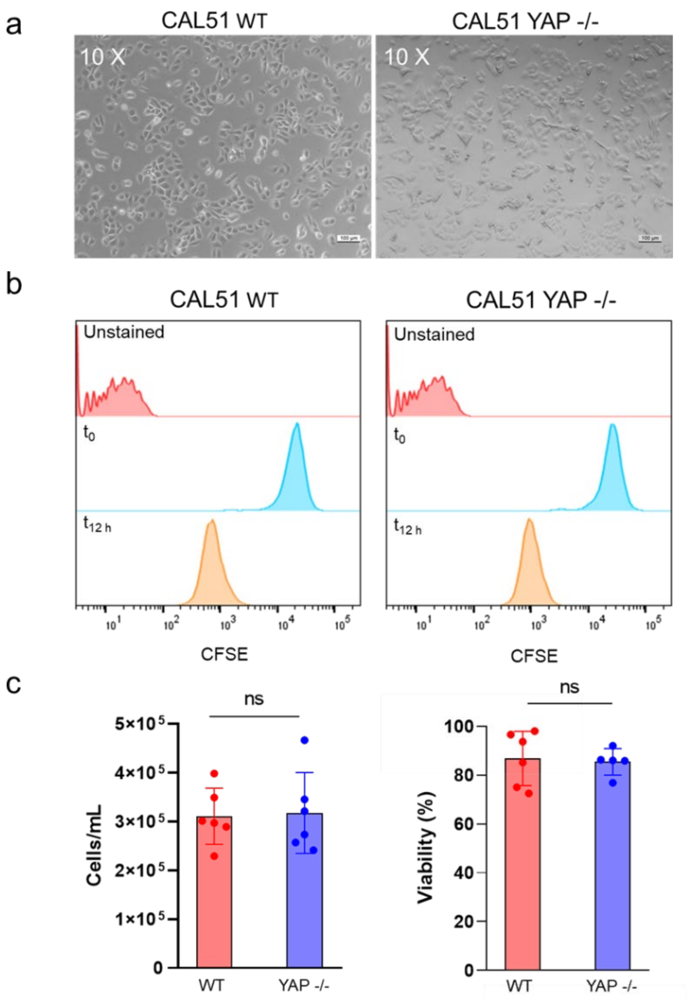

**Fig. S2.** a) Representative brightfield images of WT and YAP -/- CAL51 cells seeded onto 24-well plates 12 hours after seeding. Scale bar: 100 µm. b) Histogram of the CFSE assay for WT (left) and YAP -/- (right) cells 12 hours after seeding. Unstained cells are presented in red, cells right after seeding are presented in blue, and cells after 12 hours of seeding are presented in orange c) Cell number and viability measured *via* the trypan blue assay 12 hours after WT (red) and YAP -/- (blue) cell seeding. n = 6.

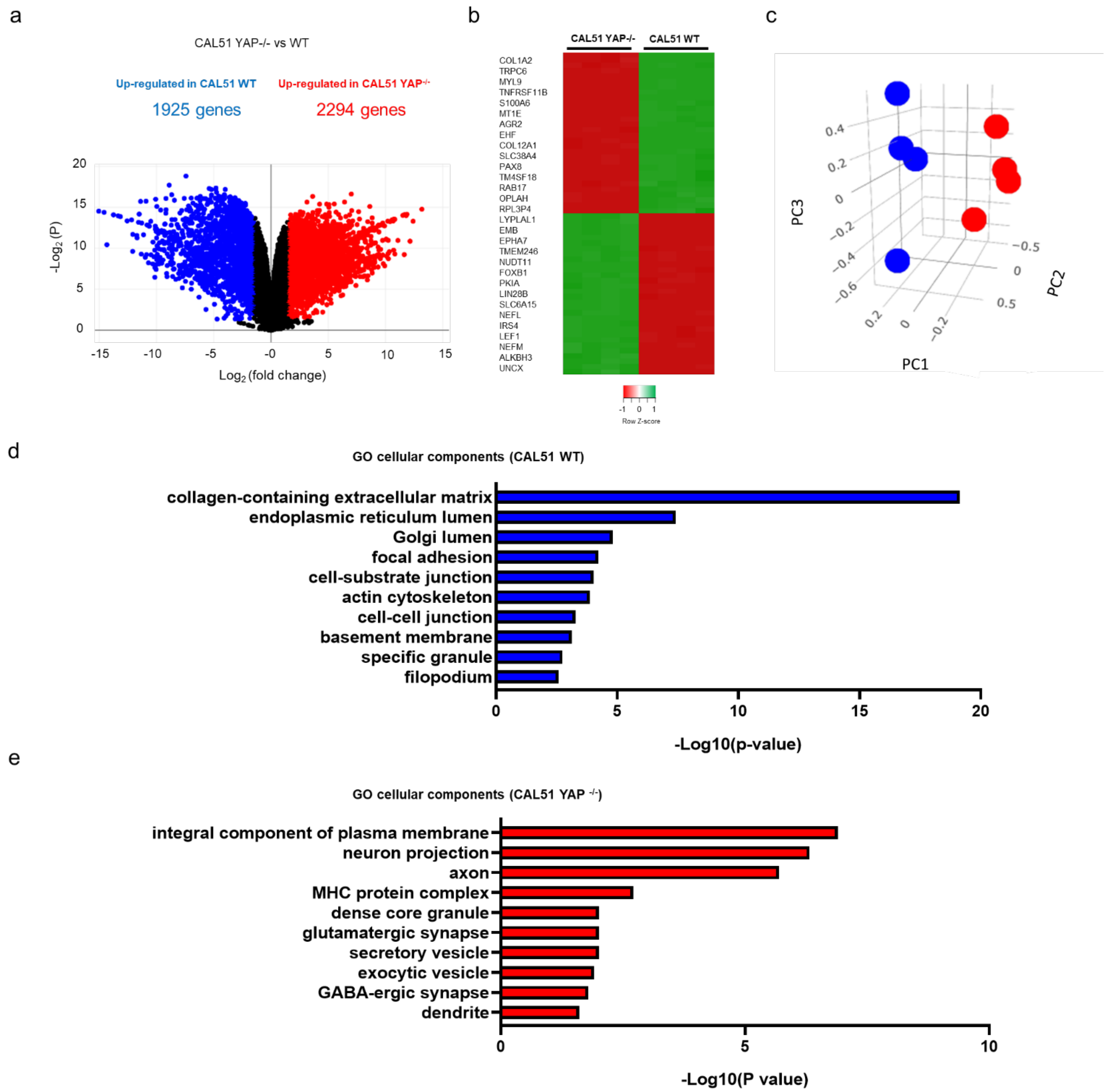

**Fig. S3.** a) Volcano plot showing differential gene expression in WT vs. YAP -/- CAL51 cells. Red points indicate significantly upregulated genes, and blue points indicate downregulated genes. N = 4 (P adj < 0.05, log2Fc > |1|). b) Heatmap of the relative gene expression of the 30 most differentially expressed genes in CAL51 WT cells compared to CAL51 YAP -/- cells (P adj < 0.05, log2Fc > |1|). c) 3D principal component analysis (PCA) of RNA-seq in CAL51 WT and CAL51 YAP -/- cells. Red dots represent a sample of WT cells, while blue dots represent a sample of YAP -/- cells. n = 4 (P adj < 0.05, log2Fc > |2|). The analysis was performed *via* Biojupies (*81*). d) Bar plot representation of common enriched cellular components obtained from the ENRICHR database (*82-84*), displaying the most significantly upregulated genes in CAL51 WT compared to CAL51 YAP -/- cells (P adj < 0.05, log2Fc > |2|). e) Bar plot representation of common enriched cellular components obtained from the ENRICHR database, displaying the most significantly upregulated genes in CAL51 YAP -/- compared to CAL51 WT cells (P adj < 0.05, log2Fc > |2|).

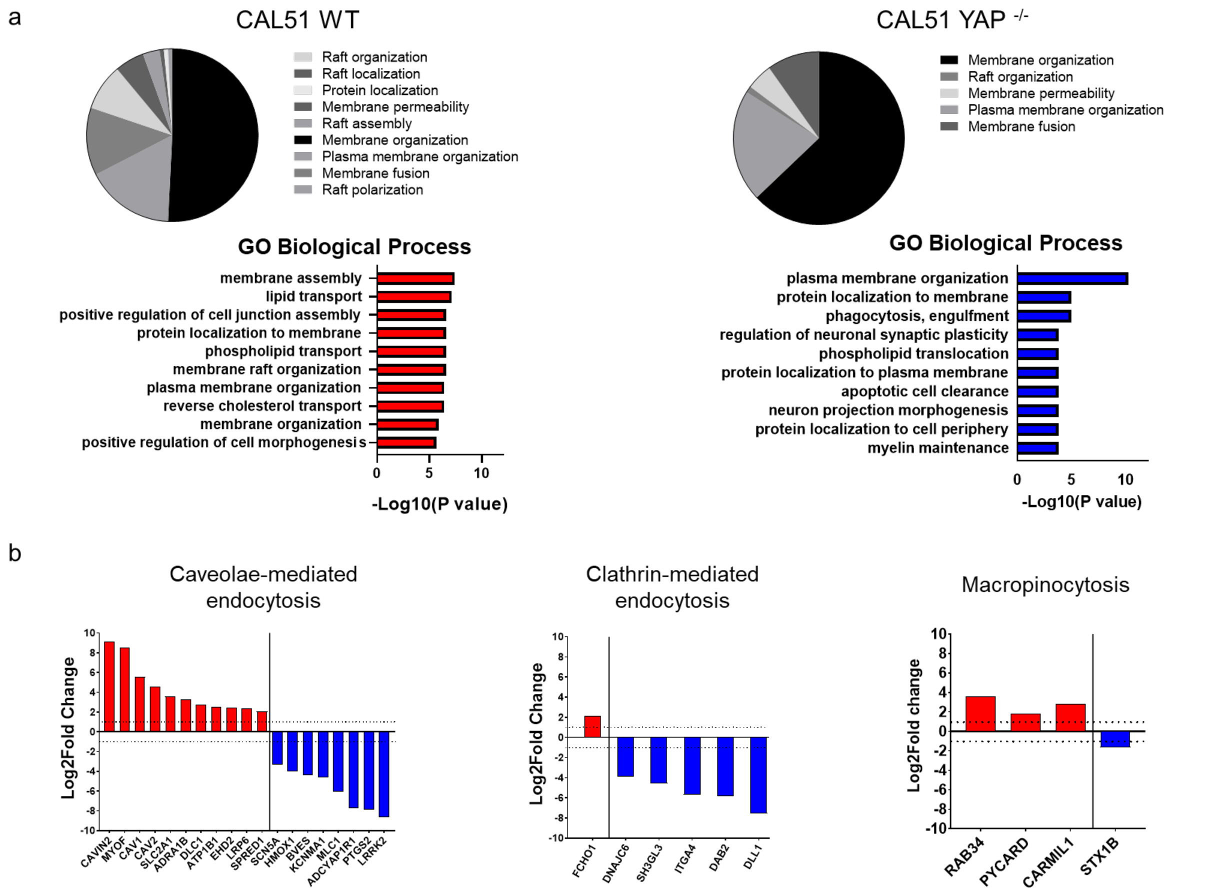

**Figure S4.** a) Pie chart of the annotations (top) and the bar plot representation (bottom) of common enriched biological processes and pathways obtained from the ENRICHR database (*82-84*), presenting the most significantly upregulated genes in WT (left) and YAP -/- (right) CAL51 cells involved in the membrane organization network (P adj < 0.05, log2Fc > |2|). b) Bar plot representations of the normalized expression of genes involved in caveolae-mediated, clathrin-mediated endocytosis and macropinocytosis found differentially regulated in YAP -/- cells as compared to WT CAL51 cells. (P adj < 0.05, log2Fc > |1|).

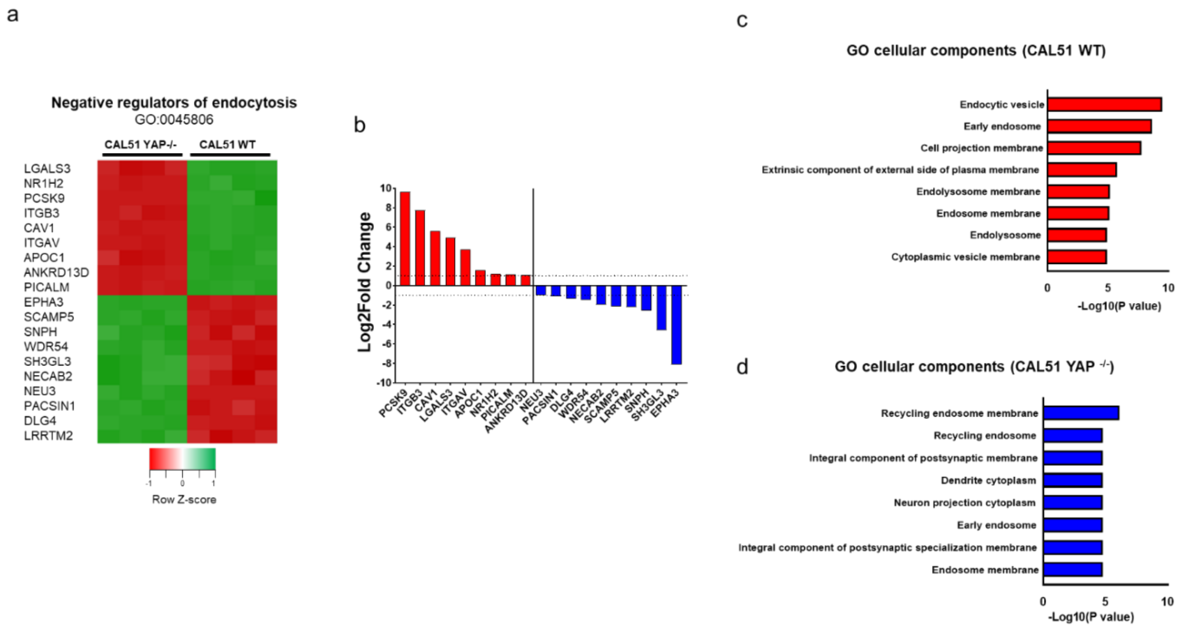

**Fig. S5.** Heatmap (a) and relative normalized expression (b) of genes involved in the negative regulation of endocytosis (GO:0045806) (P adj < 0.05, log2Fc > |1|). c) Bar plot representation of common enriched cellular components among the negative regulators of endocytosis obtained from the ENRICHR database (*82-84*), presenting the most significantly upregulated genes in WT compared to YAP -/- CAL51 cells, and d) viceversa (P adj < 0.05, log2Fc > |1|).

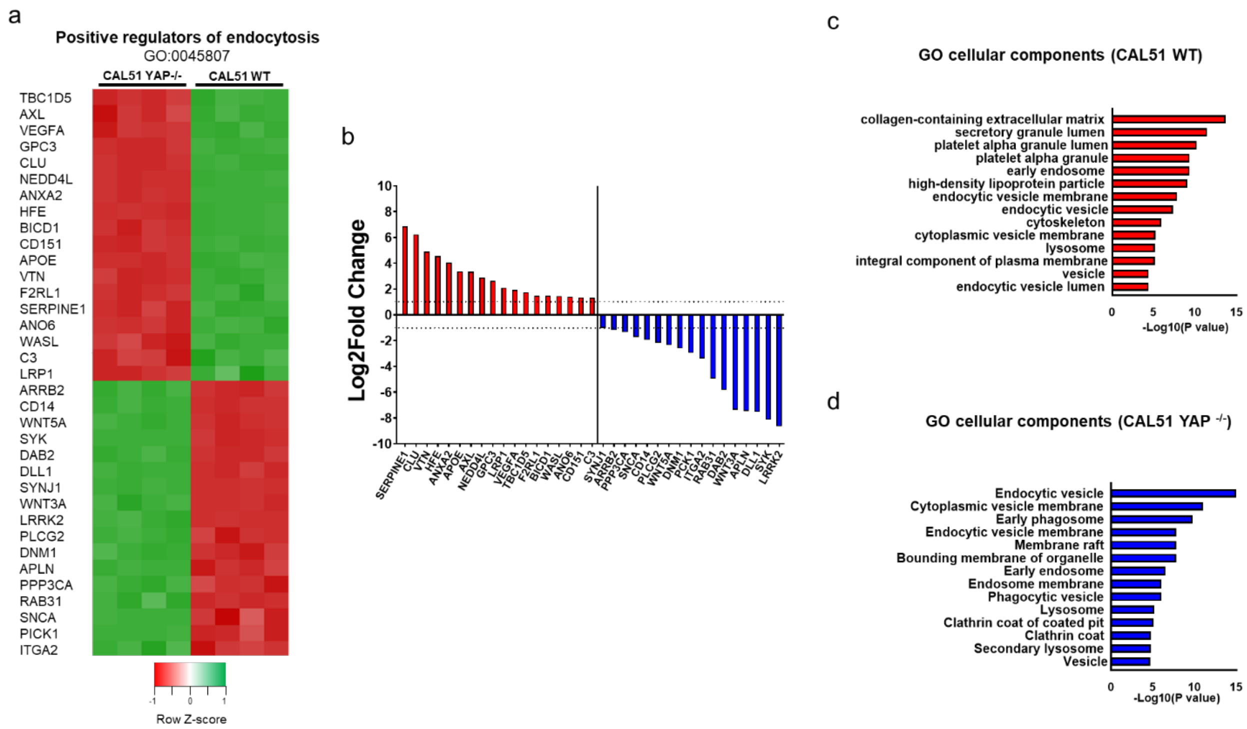

**Fig. S6.** Heatmap (a) and relative normalized expression (b) of genes involved in the positive regulation of endocytosis (GO:0045807) (P adj < 0.05, log2Fc > |1|). c) Bar plot representation of common enriched cellular components for the positive regulators of endocytosis obtained from the ENRICHR database (*82-84*), considering the most significant upregulated genes in WT compared to YAP -/- CAL51 and d) viceversa (P adj < 0.05, log2Fc > |1|).

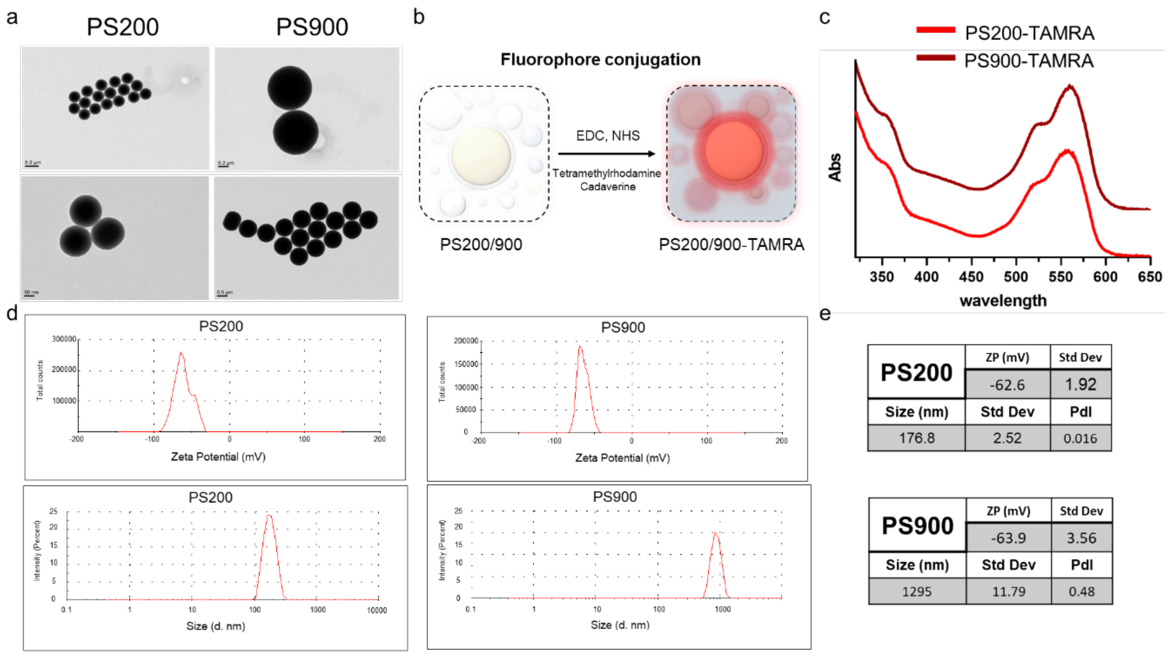

**Fig. S7.** a) Representative TEM micrographs of 200 and 900 nm polystyrene nanoparticles. Scale bars: 50, 200, and 500 nm. b) Functionalization reaction of polystyrene nanoparticles with fluorescent molecules Tetramethylrhodamine-5-carboxamide cadaverine (TAMRA cadaverine) *via* EDC chemistry. c) Normalized UV-vis absorption spectra of TAMRA-functionalized PS200 (red line) and PS900 (dark red line) nanoparticles. d) DLS graph showing the zeta potential (top) and size weighted by intensity (bottom) of PS200 and PS900 in 10 mM NaCl. e) Tables reporting the zeta potential (ZP), hydrodynamic diameter ($d_H$), and polydispersity index (PdI) of PS200 (top) and PS900 (bottom) nanoparticles.

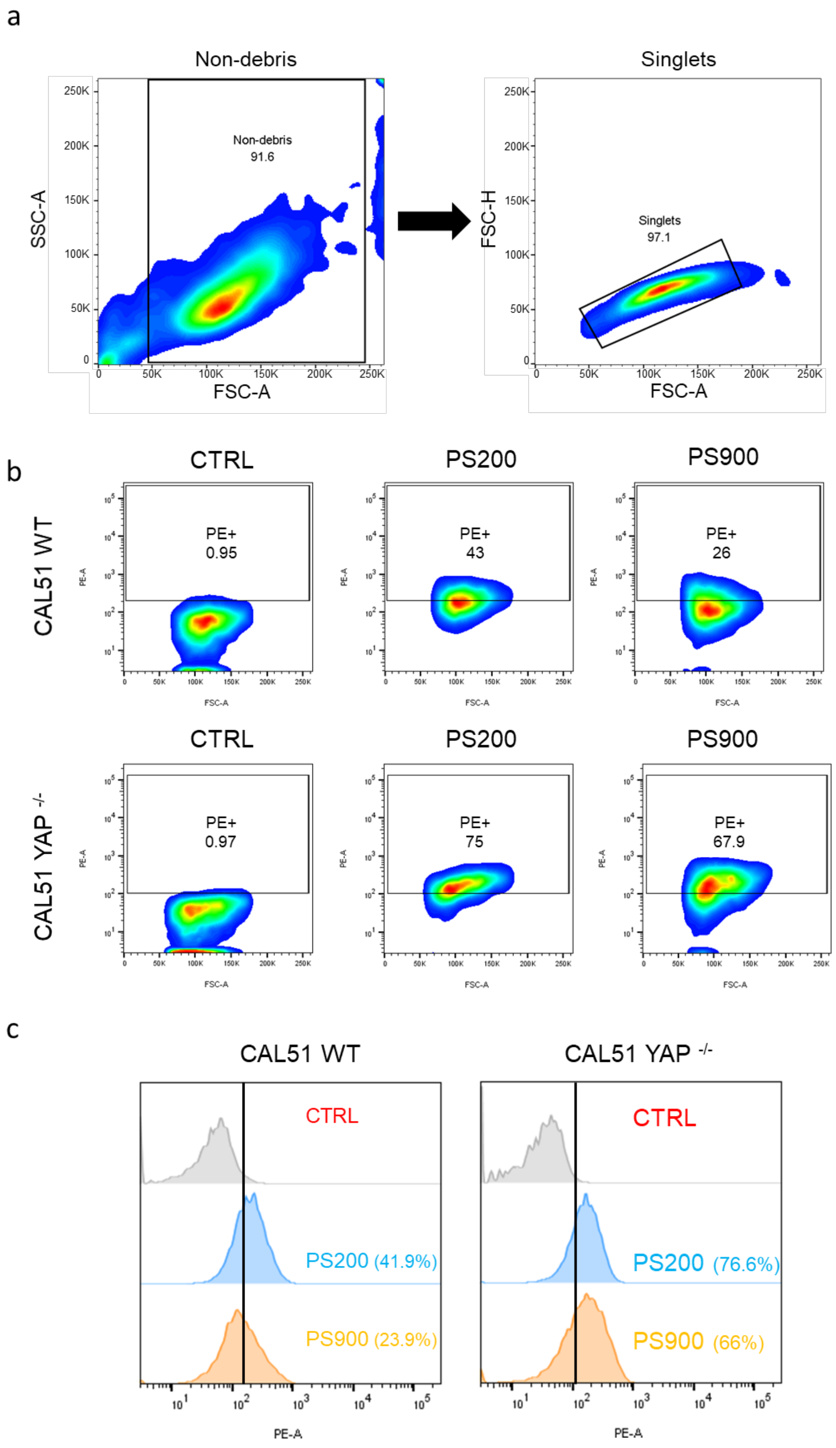

**Fig. S8.** a) Flow cytometry gating strategy for CAL51 cells. b) Representative flow cytometry plots of WT (top) and YAP -/- CAL51 cells incubated for 4 hours with PS200 and PS900. c) Representative histograms of WT (left) and YAP -/- (right) CAL51 cells incubated for 4 hours with PS200 (blue) and PS900 (orange), displaying the gating applied for PE (nanoparticle)-positive cells.

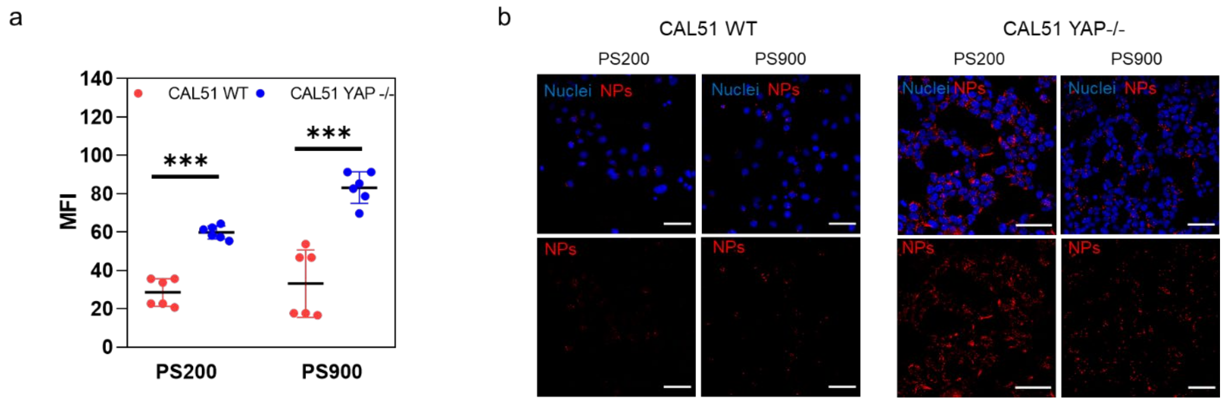

**Fig. S9.** a) Median fluorescence intensity (MFI) of a 4-hour cellular uptake of PS200 and PS900 in CAL51 WT (red) and CAL51 YAP -/- (blue) cells. Statistical analysis was performed using the two-way ANOVA with Sidak's correction for multiple comparisons. n = 6; *** indicates $p < 0.001$. b) Confocal images of WT (left) and YAP -/- (right) CAL51 cells after a 4-hour incubation with PS200 and PS900. Cells were stained with DAPI (blue). Nanoparticles are displayed in red. Scale bar: 50 µm.

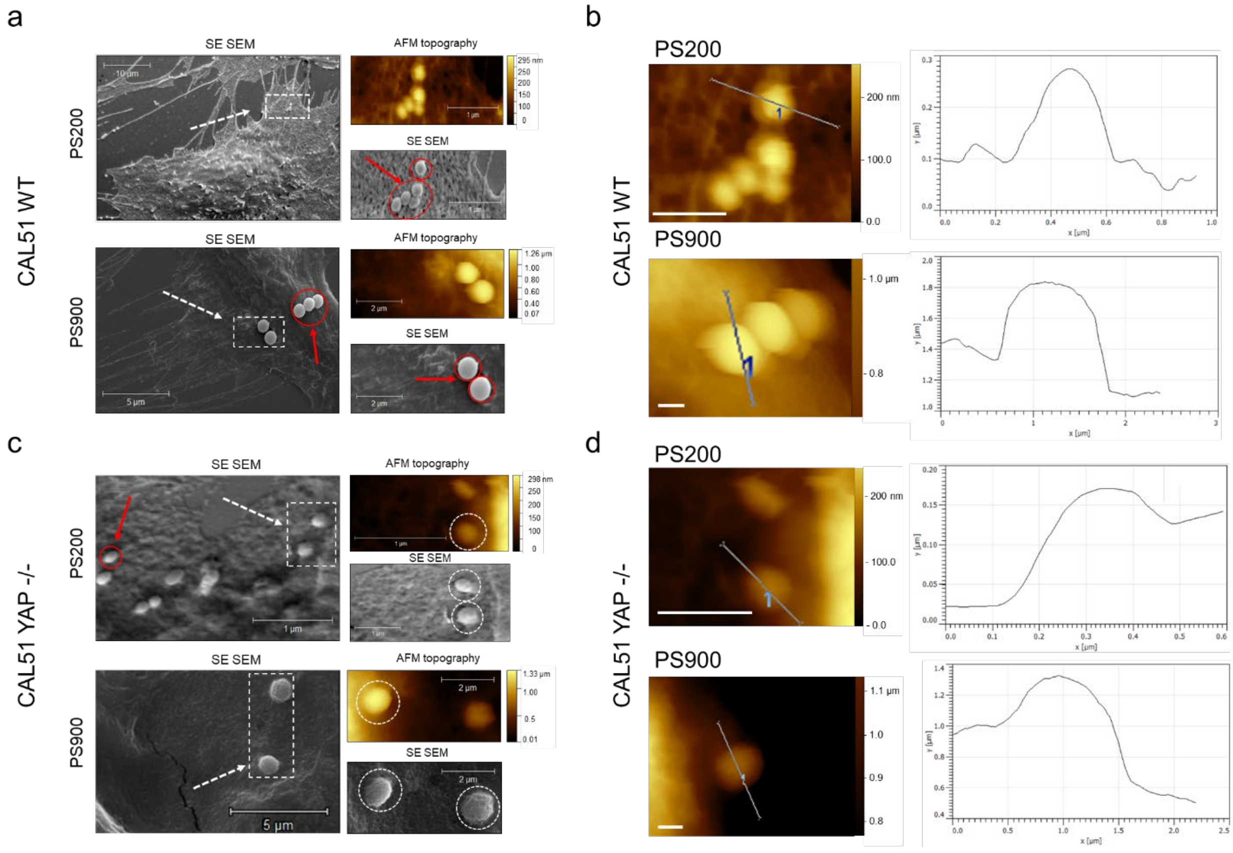

**Fig. S10.** Topographical characterization of the cell-nanoparticle membrane interaction at the membrane. a) and c) Correlative Probe and Electron Microscopy (CPEM) imaging of WT (a) and YAP -/- (c) CAL51 cells treated for 4 hours with PS200 and PS900. AFM and SEM images are presented. White dashed line boxes present magnified AFM and SEM images on the right of each micrograph. Red arrows and circles show nanoparticles bound to the cells but not embedded in the plasma membrane; white dashed arrows and circles show nanoparticles embedded in the cell membrane undergoing endocytosis. b) and d) show details of AFM images from WT (b) and YAP -/- (d) CAL51 cells incubated for 4 hours with PS200 (top) and PS900 (bottom). For each image, a plot on the right displays the height profile for the area marked by the white dashed line. All the lines where extracted using a 3 px thick line for noise reduction. The one-dimensional texture is split into waviness, reflecting the low-frequency shape components. Scale bar: 0.5 μm.

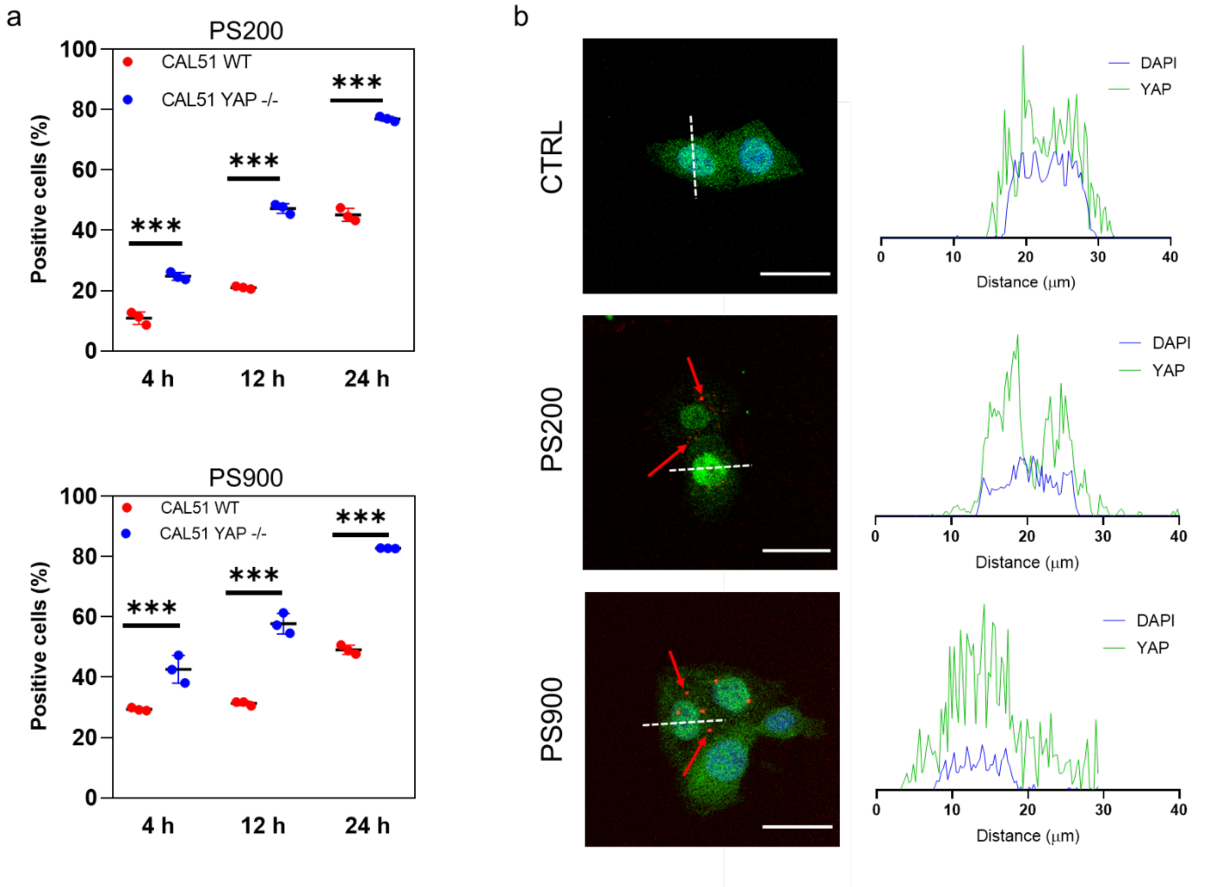

**Fig. S11.** a) 4-, 12-, and 24-hours cellular uptake of PS200 and PS900 in WT (red) and YAP -/- (blue) CAL51 cells. Statistical analysis was performed using the two-way ANOVA with Tukey's correction for multiple comparisons. n = 3; *** indicates $p < 0.001$. b) Confocal images and the intensity profile for the area marked by the white dashed line in WT CAL51 cells after 4-hour incubation with PS200 and PS900. Cells are stained with YAP (AF488) and DAPI (blue). Nanoparticles are displayed in red and pointed at by red arrows. Scale bar: 50 μm.

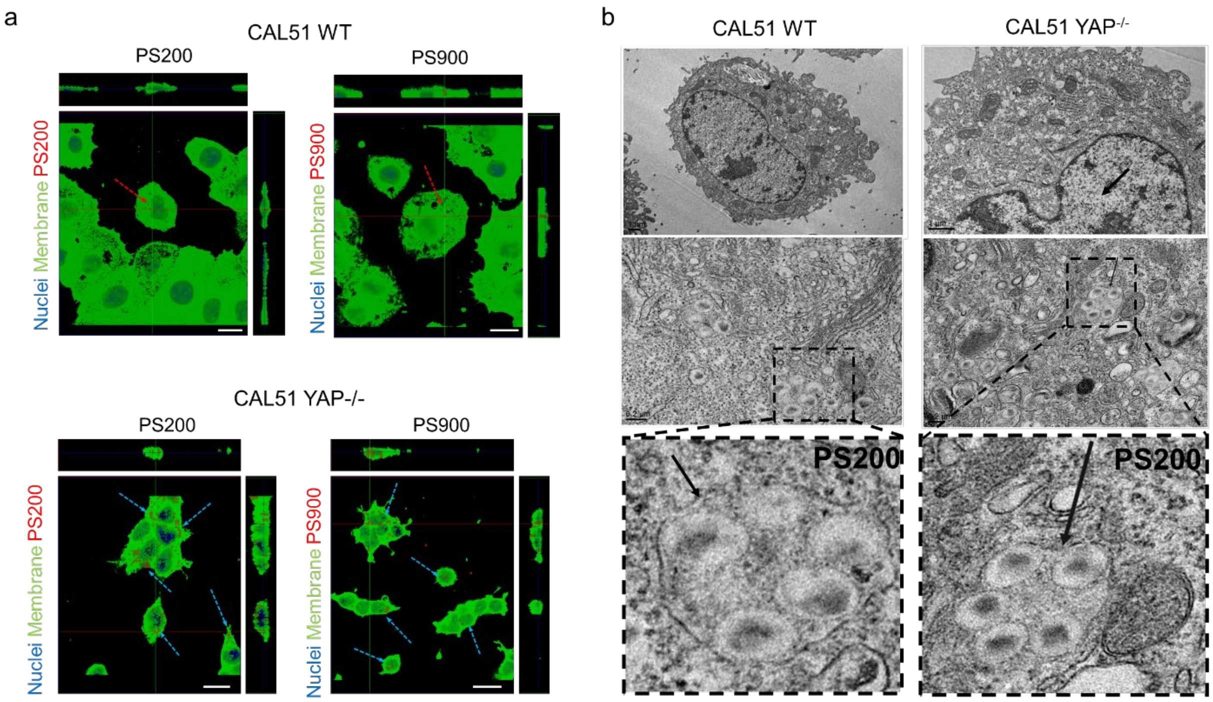

**Fig. S12.** a) Orthogonal view of z-projections of WT (top) and YAP -/- (bottom) CAL51 after 4 hours of incubation with PS200 and PS900. Cells are stained with WGA-488 (green) and DAPI (blue). Red dashed arrows indicate nanoparticles found in WT, and blue dashed arrows indicate nanoparticles found in YAP -/- CAL51. Scale bar: 20 µm. b) Thin-section TEM images of WT (top) and YAP -/- (bottom) CAL51 cells incubated with PS200 for 4 hours. Magnified images are displayed inside black dashed line boxes for each cell type. Nanoparticles are pointed at by black arrows. Scale bars: 1 and 0.2 µm.

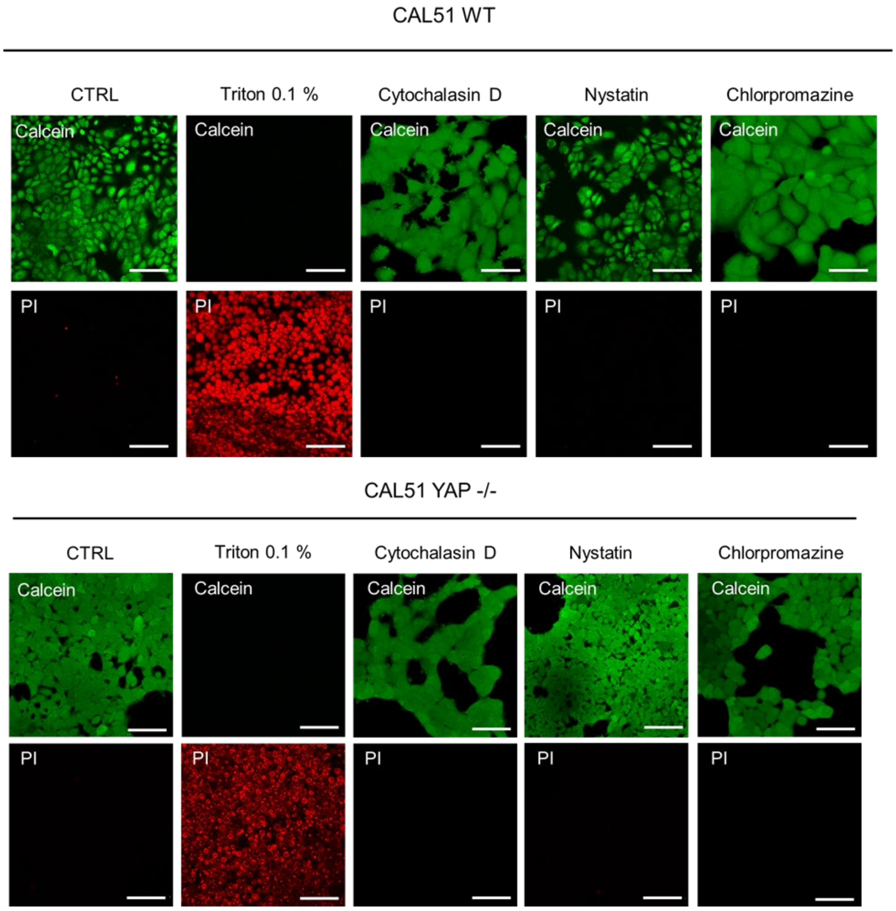

**Fig. S13.** The live/dead assay performed on WT (top) and YAP -/- (bottom) CAL51 cells treated with endocytosis inhibitors Cytochalasin D, Nystatin, and Chlorpromazine for 6 hours. As a control for cell death, cells were heated for 15 minutes with 0.1% Triton X-100 solution. Cells were excited with 555 nm laser (propidium iodide, PI, red) and 488 nm laser (calcein, green). Scale bar: 100 µm.

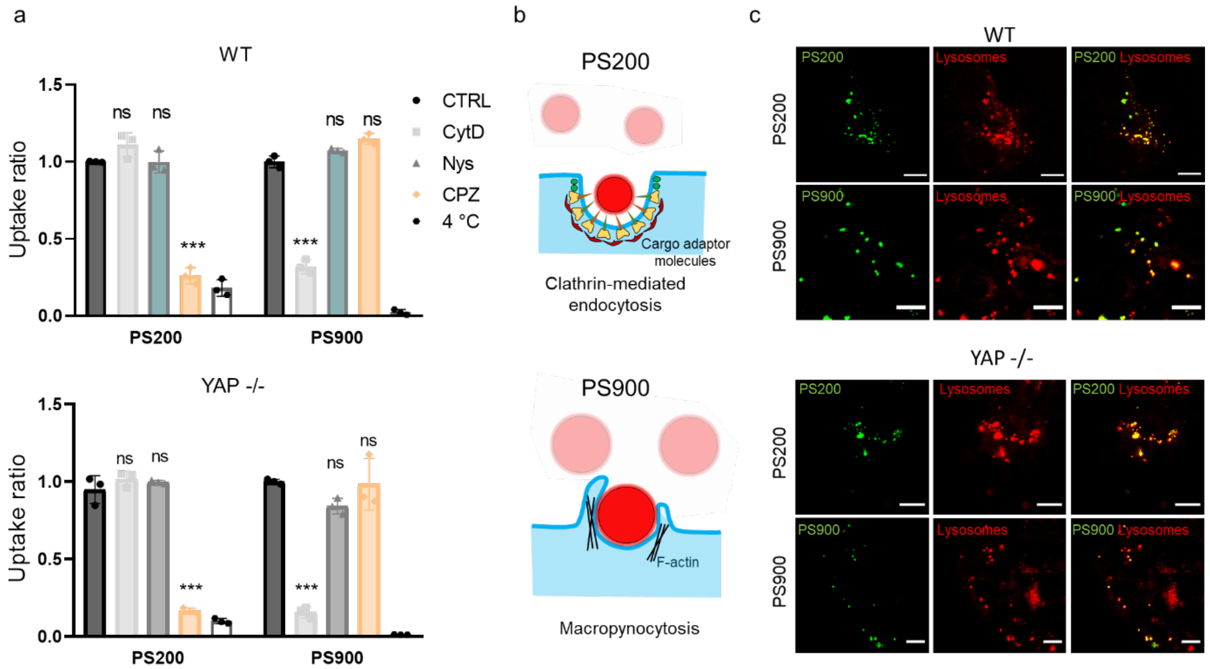

**Fig. S14. The landscape of endocytic pathways in WT and YAP -/- CAL51 cells.** a) Uptake ratios for PS200 and PS900 in WT (top) and YAP -/- (bottom) CAL51 cells upon treatment with endocytosis pathway inhibitors cytochalasin D (CytD), Nystatin (Nys), and chlorpromazine (CPZ), or incubated with nanoparticles at 4° C. Statistical analysis was performed using the two-way ANOVA with Sidak's correction for multiple comparisons. n = 3; *** indicates $p < 0.001$. b) PS200 are mainly internalized by the cells *via* clathrin-mediated endocytosis (top), while PS900 undergo macropinocytosis (bottom). c) Confocal images of the intracellular localization of PS200 (top) and PS900 (bottom) in CAL51 WT (left) and CAL51 YAP -/- (right) cells 8 hours after 4-hour incubation with the nanoparticles. Cells are stained with Lysotracker. Nanoparticles are displayed in green. Scale bar: 10 µm

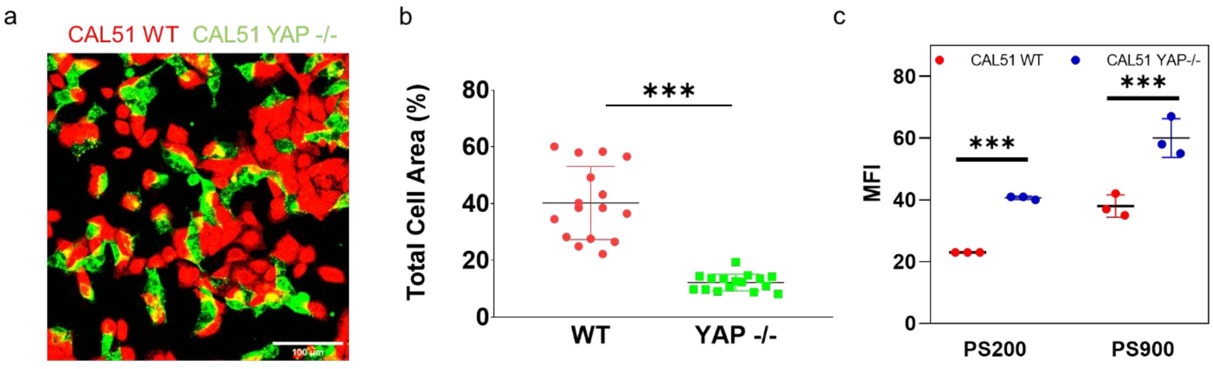

**Fig. S15.** a) Confocal image of the WT and YAP -/- CAL51 co-culture. WT cells are stained with PKH-red, and YAP -/- cells are stained with PKH-green. Scale bar: 100 µm. b) Area of WT and YAP -/- CAL51 cells in a co-culture calculated based on the total membrane area of the cells. Statistical analysis was performed by unpaired t-test using Welch's correction. n > 10; *** indicates $p < 0.001$. c) Median fluorescence intensity (MFI) of a 4-hour cellular uptake of PS200 and PS900 for WT (red) and YAP -/- (blue) CAL51 cells in co-culture. Statistical analysis was performed using the two-way ANOVA with Sidak's correction for multiple comparisons. n = 3; *** indicates $p < 0.001$.

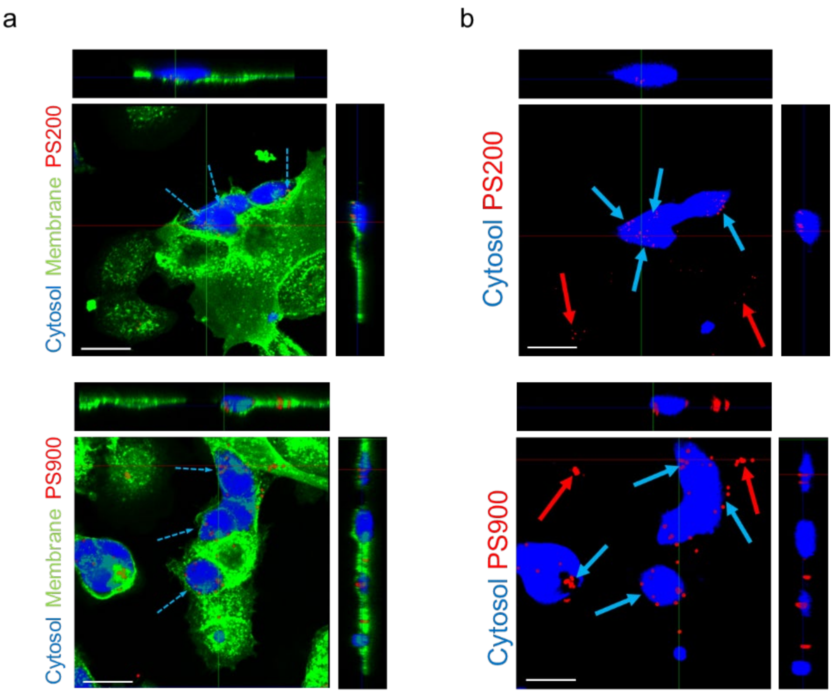

**Fig. S16** a) Orthogonal view of z-projections of the co-culture after 4 hours of incubation with PS200 (top) and PS900 (bottom). Blue dashed arrows indicate the particles found in YAP -/- CAL51 cells. YAP -/- cells are stained with 7-amino-4-chloromethylcoumarin (blue) and whole cell population with WGA-488 (green). Scale bar: 20 µm. b) Orthogonal view of the Z-projections of the co-culture after 4 hours of incubation with PS200 (left) and PS900 (right). Blue arrows point at the nanoparticles found in YAP -/- cells, while red arrows point to the particles found in WT CAL51 cells. YAP -/- CAL51cells were stained with 7-amino-4-chloromethylcoumarin (blue). Scale bar: 20 µm.

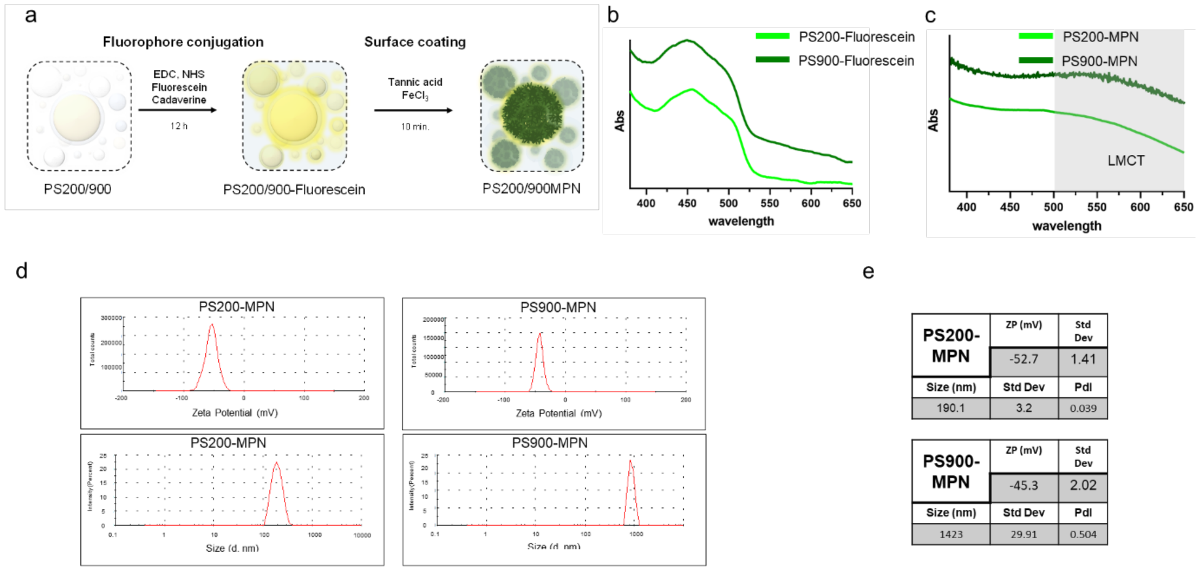

**Fig. S17.** a) Functionalization reaction of polystyrene nanoparticles with fluorescent molecules Tetramethylrhodamine-5-carboxamide cadaverine (fluorescein cadaverine) *via* EDC chemistry and the subsequent metal phenolic network (MPN) formation using tannic acid (TA) and iron chloride ($FeCl_3$). b) Normalized UV-vis absorption spectra of fluorescein-functionalized PS200 (green line) and PS900 (dark green line) nanoparticles. c) Normalized UV-vis absorption spectra of MPN-coated fluorescein-functionalized PS200 (green line) and PS900 (dark green line) nanoparticles. The shadowed area indicates the absorbance region of the ligand-to-metal charge transfer (LMCT) of the particles coated with MPN. d) DLS graph showing the zeta potential (top) and size weighted by intensity (bottom) of PS200 and PS900 in 10 mM NaCl. e) Tables reporting the zeta potential (ZP), hydrodynamic diameter ($d_H$), and polydispersity index (PdI) of PS200-MPN (top) and PS900-MPN (bottom) nanoparticles.

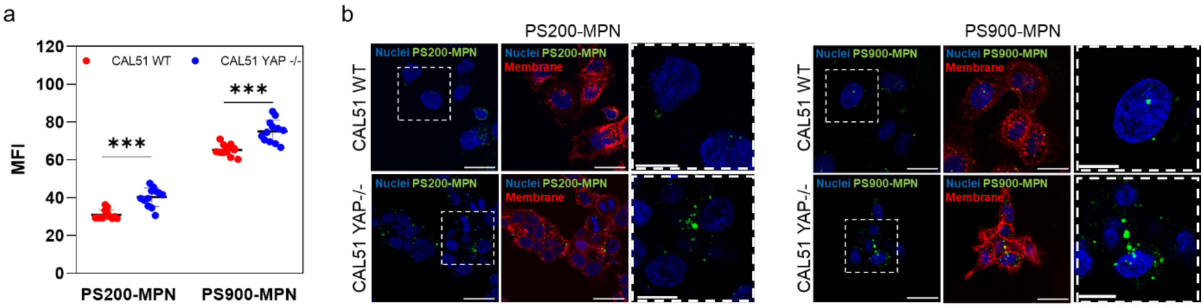

**Fig. S18.** b) Median fluorescence intensity (MFI) of a 4-hour cellular uptake of PS200-MPN and PS900-MPN for WT (red) and YAP -/- (blue) CAL51cells. Statistical analysis was performed using the two-way ANOVA with Tukey's correction for multiple comparisons. n = 12; *** indicates p < 0.001. b) Confocal images of WT (left) and YAP -/- (right) CAL51 cells after 4-hour incubation with PS200-MPN and PS900-MPN. Cells are stained with WGA-647 (red) and DAPI (blue). Nanoparticles are displayed in red. Magnified images are presented inside the white dashed line boxes on the right relative to the highlighted sections on the left. Scale bars: 20 and 10 µm.

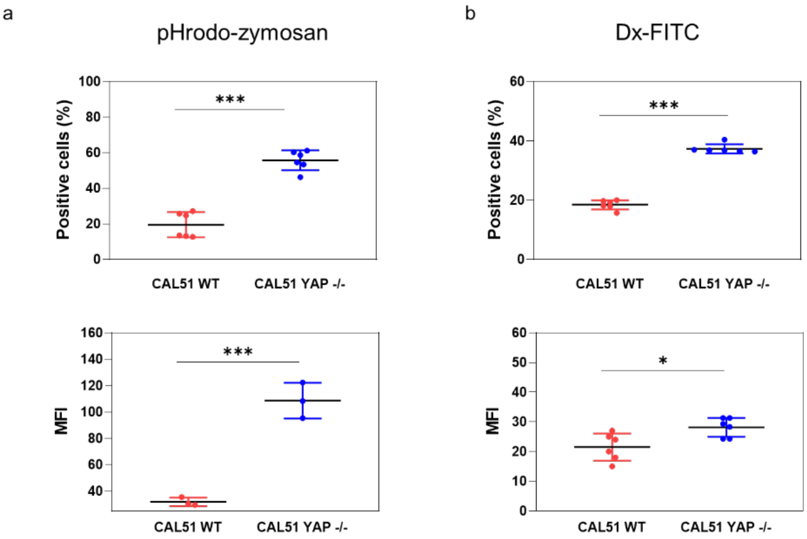

**Fig. S19.** a) 4-hour cellular uptake of pHrodo-zymosan expressed as a percentage of positive cells (top) and median fluorescence intensity (MFI) for WT (red) and YAP +/+ (blue) CAL51 cells. Statistical analysis was performed using the unpaired t-test with Welch's correction. n = 6 and n = 3; *** indicates p < 0.001. b) 4-hour cellular uptake of Dx-FITC expressed as a percentage of positive cells (top) and MFI for WT (red) and YAP +/+ (blue) CAL51 cells. Statistical analysis was performed using the unpaired t-test with Welch's correction. n = 6; * indicates p < 0.05; *** indicates p < 0.001.

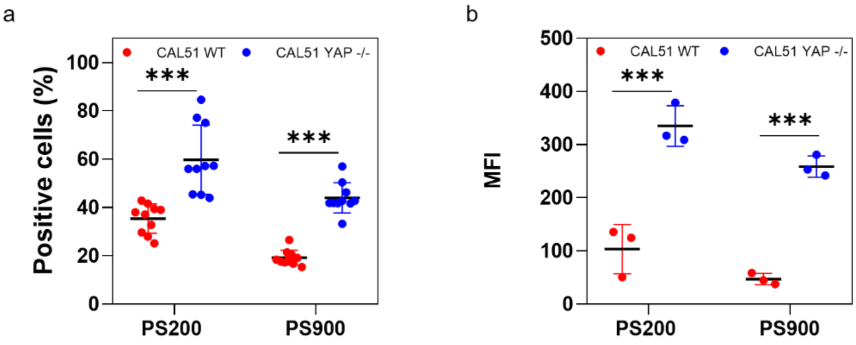

**Fig. S20.** a) 4-hour cellular uptake of PS200 and PS900 in a serum-free medium for CAL51 WT (red) and CAL51 YAP -/- (blue) cells. Statistical analysis was performed using the two-way ANOVA with Sidak's correction for multiple comparisons. n = 9; *** indicates p < 0.001. b) Median fluorescence intensity (MFI) of 4-hour cellular uptake of PS200 and PS900 in a serum-free medium for CAL51 WT (red)

and CAL51 YAP +/+ (blue) cells. Statistical analysis was performed using the two-way ANOVA with Tukey's correction for multiple comparisons. n = 3; *** indicates $p < 0.001$.

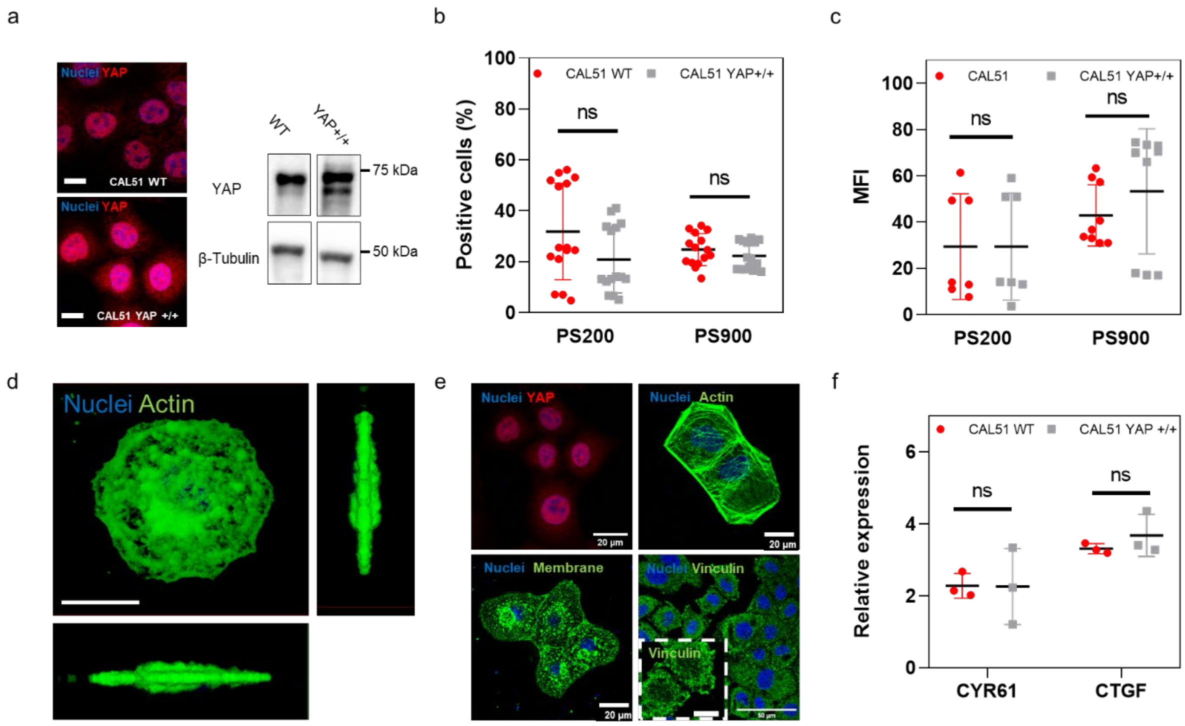

**Fig. S21.** a) Confocal images of CAL51 WT cells (top) and CAL51 YAP +/+ cells, transfected with YAP (right) (*85*). Cells are stained for YAP (AF555) and DAPI. Scale bar: 10 µm. A western blot on the right shows YAP levels in CAL51 WT and YAP +/+ cells. b) 4-hour cellular uptake of PS200 and PS900 in CAL51 WT (red) and CAL51 YAP +/+ (blue) cells. Statistical analysis was performed using the two-way ANOVA with Tukey's correction for multiple comparisons. n = 15; ns indicates non-significant. c) Median fluorescence intensity (MFI) of a 4-hour cellular uptake of PS200 and PS900 for CAL51 WT (red) and CAL51 YAP +/+ (blue) cells. Statistical analysis was performed using the two-way ANOVA with Tukey's correction for multiple comparisons. n = 6; ns indicates non-significant. d) 3D reconstruction of CAL51 YAP +/+ cells. The top and lateral views are presented. The cells are stained with DAPI (blue) and WGA-488 (green). Scale bar: 20 µm. e) Confocal images of CAL51 YAP +/+ cells stained for DAPI (blue), YAP (AF555, red, top left), actin (AF488, green, top right), membrane (WGA-647, red), and vinculin (AF488, green, bottom right). The bottom right panel shows vinculin staining in a magnified image inside the white dashed line box. Scale bars: 20 and 10 µm. f) Quantitative RT-PCR analysis of CYR61 and CTGF in CAL51 WT (red) and CAL51 YAP +/+ (grey) cells. Statistical analysis was performed using the unpaired t-test with Welch's correction. n = 3; ns indicates non-significant.

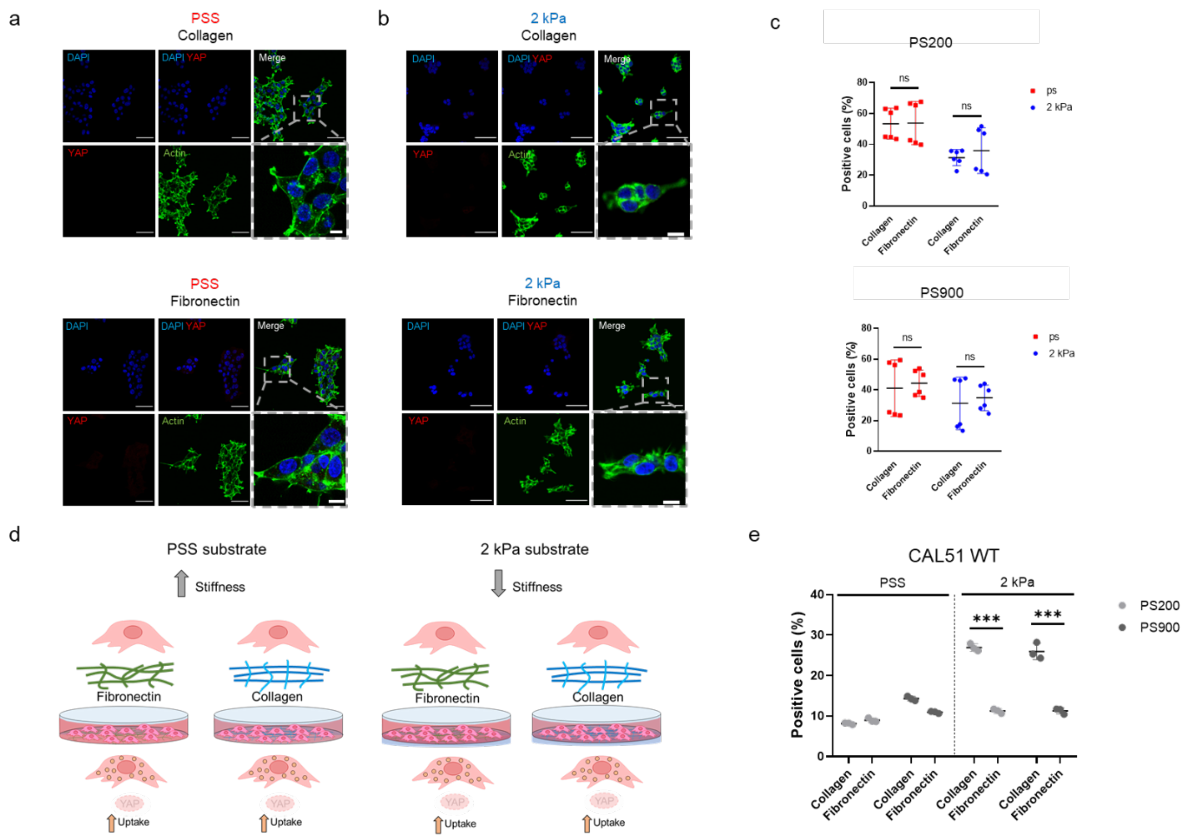

**Fig. S22.** a) Confocal images of CAL51 YAP -/- cells grown on a standard polystyrene substrate coated with collagen (top) or fibronectin (bottom). Cells are stained with DAPI (blue) and Pha-488 (green), as well as for YAP (red). Magnified images are presented inside the gray dashed line boxes. Scale bars: 50 and 10 µm. b) Confocal images of CAL51 YAP -/- cells grown on 2 kPa soft substrate coated with collagen (top) or fibronectin (bottom). Cells are stained with DAPI (blue) and Pha-488 (green), as well as for YAP (red). Magnified images are presented inside the gray dashed line boxes. Scale bars: 50 and 10 µm. c) 4-hour cellular uptake of PS200 (top) and PS900 (bottom) for CAL51 YAP -/- cells grown on polystyrene (red) or soft 2 kPa (blue) substrate coated with collagen or fibronectin. Statistical analysis was performed using the two-way ANOVA with Sidak's correction for multiple comparisons. n = 3; ns is non-significant. d) When CAL51 YAP -/- cells are grown on polystyrene or soft substrates, nanoparticle uptake remains consistently higher compared to CAL5 WT cells, regardless of the coating material used for the substrate. e) Comparison of the 4-hours cellular uptake of PS200 (light grey) and PS900 (dark grey) by CAL51 WT cells grown on PSS (left) or 2 kPa (right) substrates coated with collagen or fibronectin. Statistical analysis was performed using the two-way ANOVA with Sidak's correction for multiple comparisons. n = 3; *** indicates $p < 0.001$.

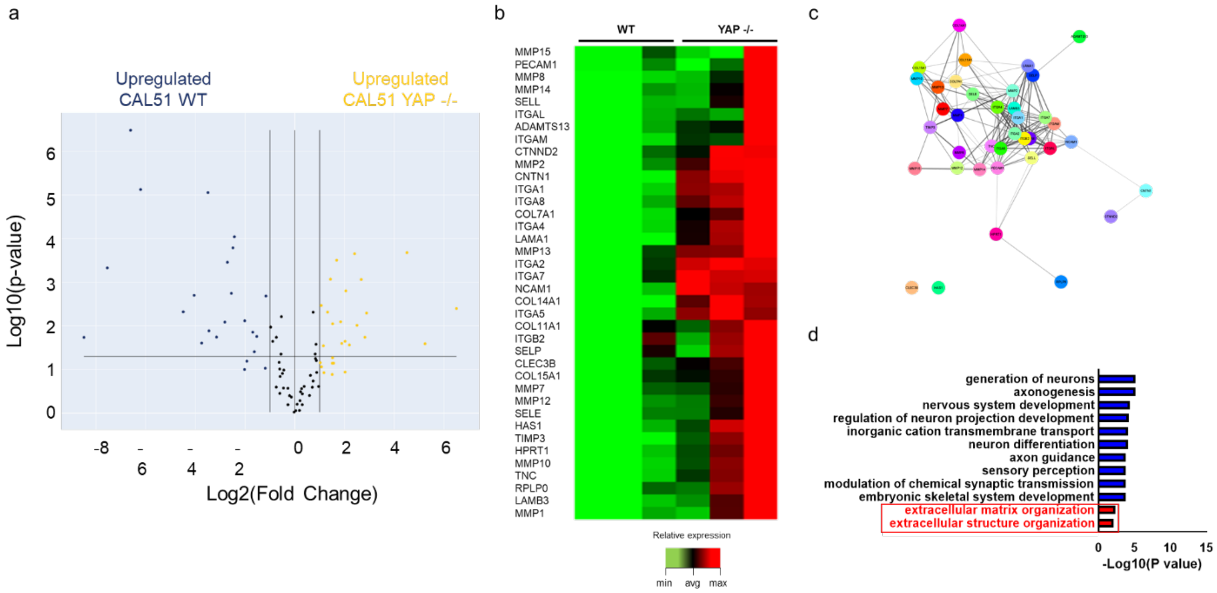

**Fig. S23.** a) Volcano plot representation of the differentially regulated genes in CAL51 WT vs. CAL51 YAP -/- cells. Blue points indicate significantly upregulated genes in CAL51 WT, and yellow points indicate upregulated genes in CAL51 YAP -/-. n = 12 (P adj < 0.05, log2Fc > |1|). b) RT$^2$-profiler PCR array for the ECM and adhesion molecules overexpressed in CAL51 YAP -/- compared to CAL51 WT (P adj < 0.05, log2Fc > |1|). c) STRING PPI network of the differently expressed ECM proteins in CAL51 YAP -/- obtained from Cytoscape (P adj < 0.05, log2Fc > |2|, confidence cutoff 0.4). d) Bar plot representation of common enriched biological processes and pathways obtained from the ENRICHR database (*82-84*), presenting the most significantly upregulated genes in CAL51 YAP -/- compared to CAL51 WT. Annotations directly related to ECM organization are highlighted in the red box (P adj < 0.05, log2Fc > |2|).

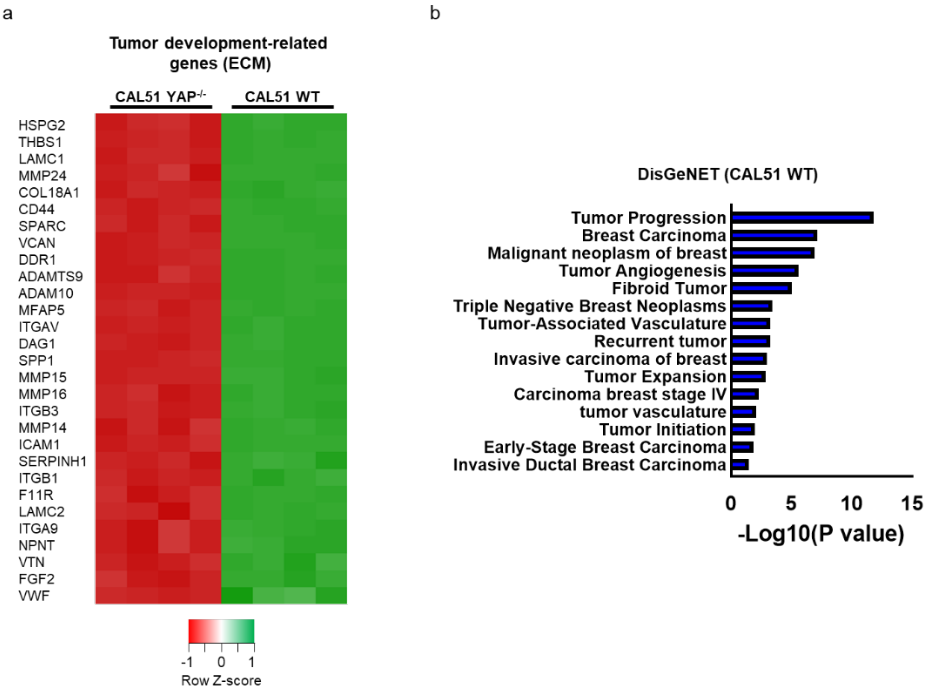

**Fig. S24.** a) Heatmap of the relative expression of genes related to ECM components and involved in tumor development (P adj < 0.05, log2Fc > |2|). b) Bar plot representation of common enriched pathways obtained from the DisGeNET database, presenting the most significantly upregulated genes in CAL51 WT compared to CAL51 YAP -/- cells (P adj < 0.05, log2Fc > |2|).

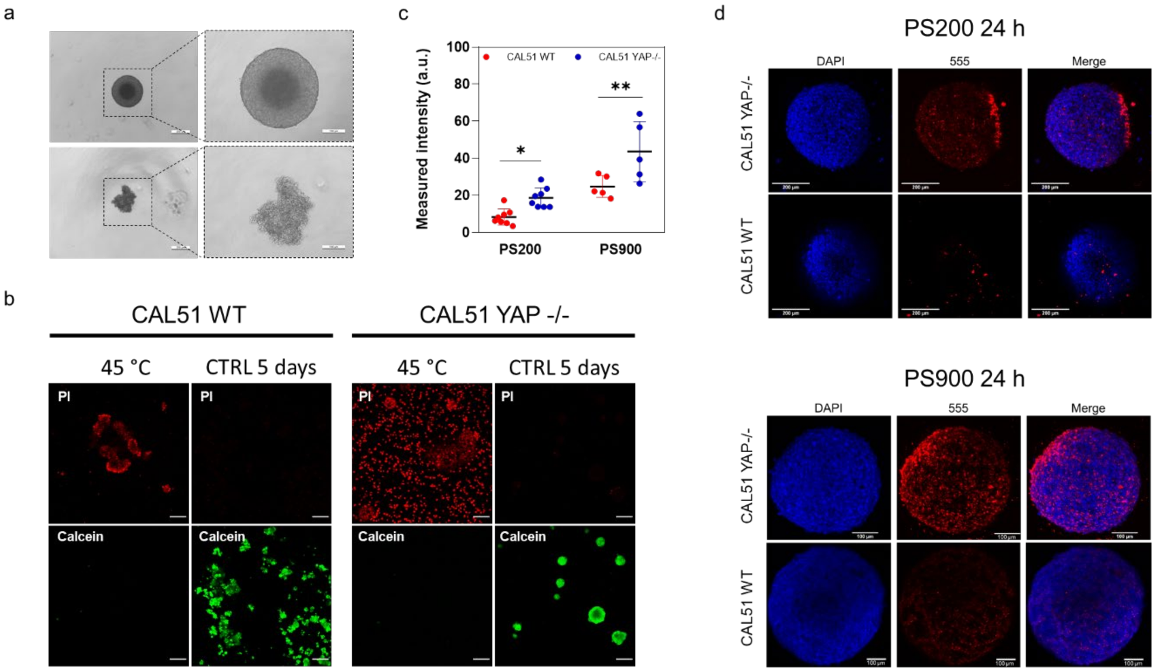

**Fig. S25.** a) Representative brightfield images of CAL51 WT and CAL51 YAP -/- spheroids seeded onto 96-well round (U) bottom plates 5 days after seeding. Magnified images are presented inside the black dashed line boxes. Scale bars: 200 and 100 µm. b) The live/dead assay performed on CAL51 WT and CAL51 YAP -/- cells. As a control for cell death, spheroids were heated for 10 minutes at 45 °C. Cells were excited with 555 nm laser (propidium iodide, PI, red) and 488 nm laser (calcein, green). c) Nanoparticle intensity per cell after 4-hour incubation of CAL51 WT and YAP -/- cells with PS200 and PS900. Statistical analysis was performed using the two-way ANOVA with Sidak's correction for multiple comparisons. $n > 5$; * indicates $p < 0.05$, ** indicates $p < 0.01$. d) Confocal images of CAL51 YAP -/- and CAL51 WT spheroids incubated with PS200 and PS900 for 24 hours. Cells are stained with DAPI (blue). Nanoparticles are displayed in red. Scale bars: 100 and 200 µm.

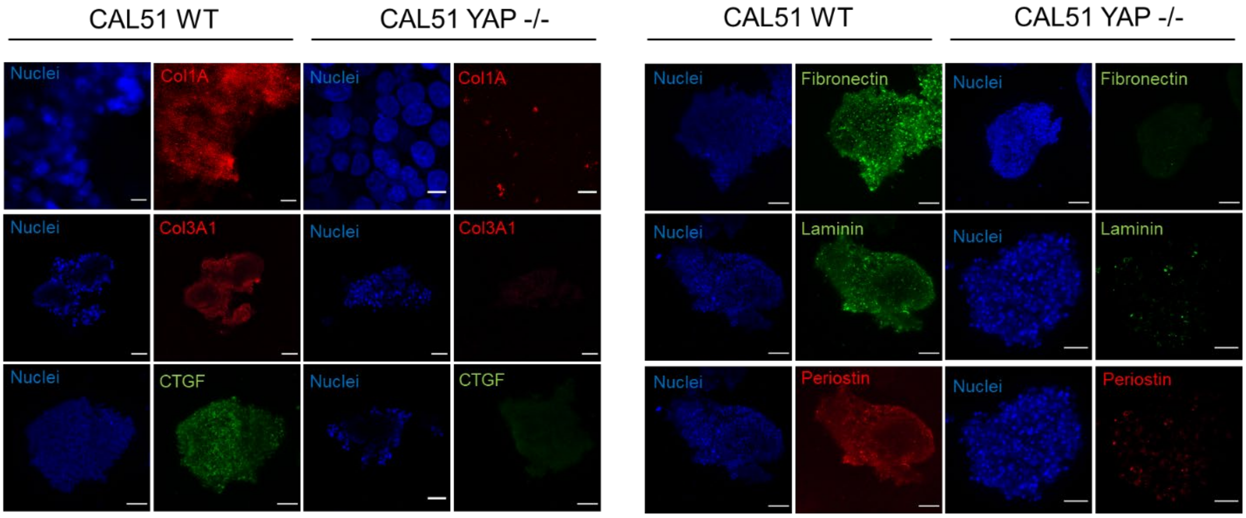

**Fig. S26.** Confocal images of ECM components for the CAL51 WT (left) and YAP -/- (right) spheroids. Collagen type 1 alpha (Col1A), collagen type III alpha 1 (Col3A1), connective tissue growth factor (CTGF), and periostin are stained with II-antibody labeled with AF-555 (red); fibronectin and laminin are stained with II-antibody labeled with AF-488 (green). Scale bar: 100 µm.

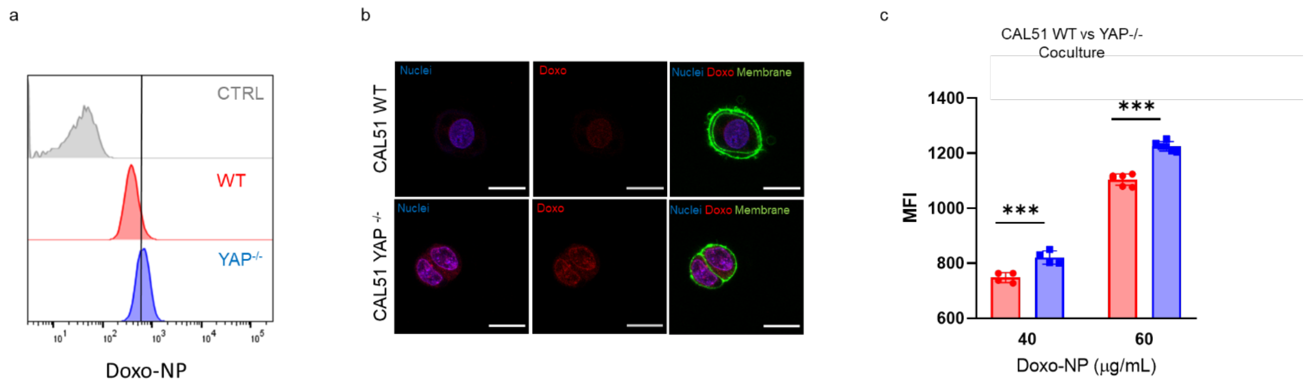

**Fig. S27.** a) Representative histograms of CAL51 WT (red) and YAP -/- (blue) cells incubated for 4 hours with Doxo-NP. b) Confocal images of CAL51 WT (top) and CAL51 YAP -/- (bottom) cells after 8-hour incubation with Doxo-NP. Cells are stained with WGA-488 (green) and DAPI (blue). Nanoparticles are displayed in red. Scale bar: 20 µm. c) Median fluorescence intensity (MFI) of 4-hour cellular uptake of Doxo-NP at different concentrations (40 and 60 µg/mL) for CAL51 WT (red) and CAL51 YAP -/- (blue) cells in co-culture. Statistical analysis was performed using the two-way ANOVA with Sidak's correction for multiple comparisons. n = 6; *** indicates $p < 0.001$.

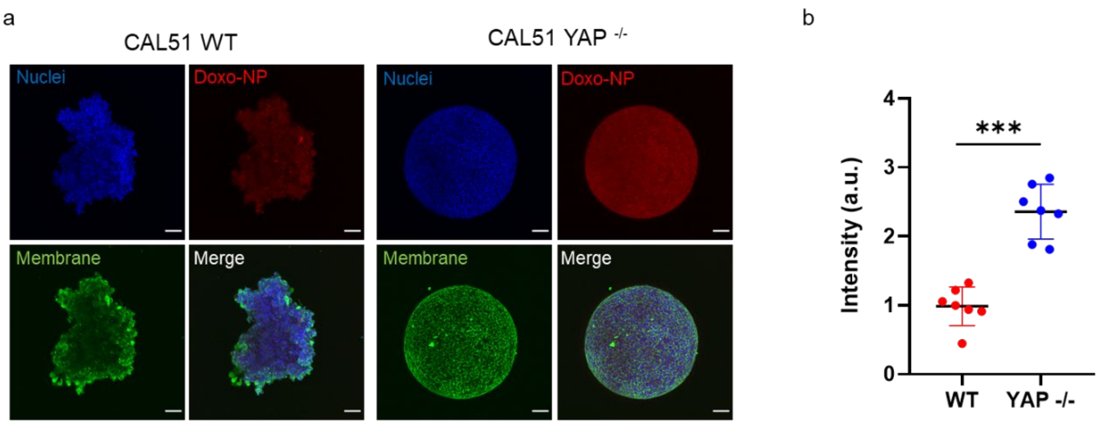

**Fig. S28.** a) Confocal images of spheroids from CAL51 WT (left) and YAP -/- (right) cells incubated for 4 hours with Doxo-NP. Cells are stained with WGA-488 (green) and DAPI (blue). Nanoparticles are displayed in red. Scale bar: 100 µm. b) Nanoparticle intensity per spheroid after 4-hour incubation of CAL51 WT (red) and YAP -/- (blue) spheroids with Doxo-NP. Statistical analysis was performed using the unpaired t-test with Welch's correction. n = 7; *** indicates $p < 0.001$.

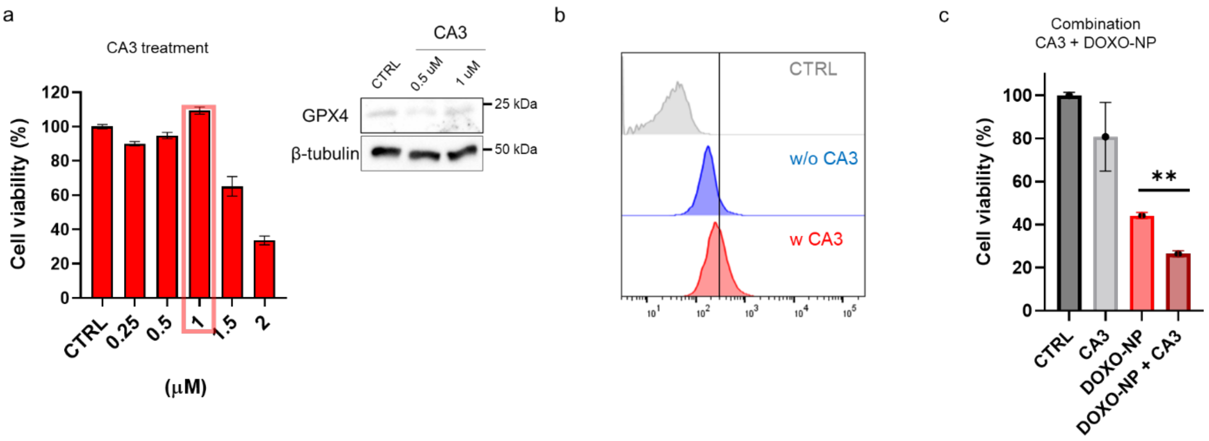

**Fig. S29.** a) Presto Blue viability assay on CAL51 WT cells treated with different concentrations of CA3 (0.25/0.5/1/1.5/2 µM) for 12 hours. The red box indicates the concentration chosen for the follow up experiments. n = 8. On the right, western blot showing the levels of ferroptosis marker GPX4 in CAL51 WT untreated (CTRL) or treated with 0.5 and 1 µM CA3 inhibitor for 12 hours. β-tubulin was used for protein loading normalization. b) Representative histograms of CAL51 WT cells untreated (blue) or treated (red) with 1 µM CA3 for 12 hours and incubated for 4 hours with Doxo-NP. c) Presto Blue viability assay on CAL51 WT cells untreated (red) or treated for 12 hours with 1 µM CA3 and subsequently incubated for 4 hours with Doxo-NP. Cell viability was assessed 48 hours post-treatment and normalized to untreated cells (control). Statistical analysis was performed using one-way ANOVA with Tukey's correction for multiple comparisons. n = 8; ** indicates $p < 0.01$.

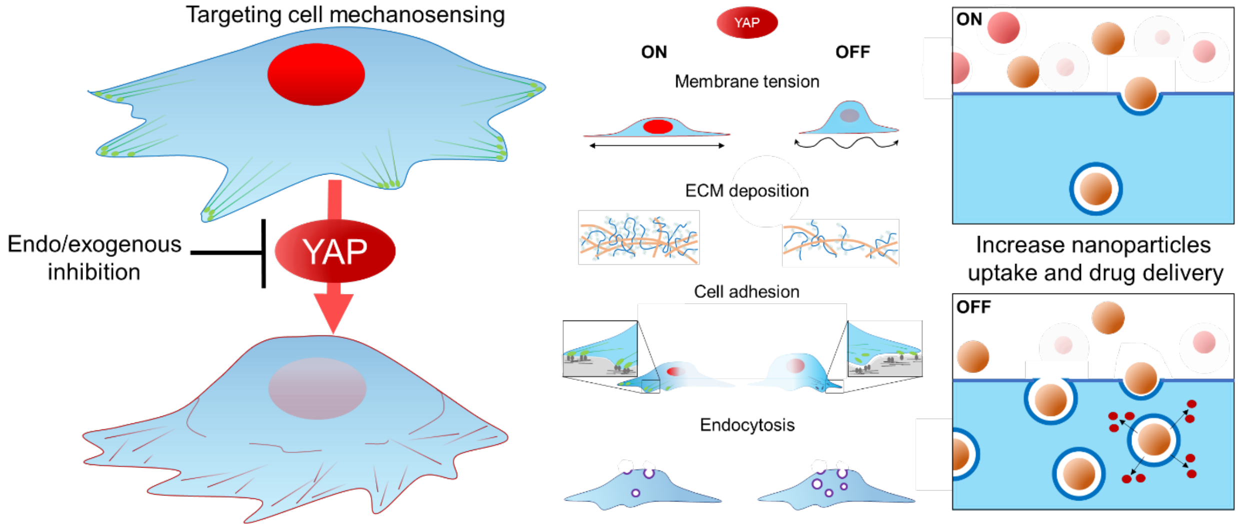

**Fig. S30.** Genetic and pharmacological inhibition of YAP in TNBC cells affects the organization of plasma membrane, reduces the ECM deposition, impacts their adhesion ability and increase the endocytosis rate. Ultimately, these changes contribute cooperatively to promote the delivery of nanomedicines to cancer cells and improve the therapeutic efficiency.

**Supplementary Table 1.** List of used immunofluorescence antibodies and their relative dilution.

| Antibody | Dilution | Buffer | Reference | Supplier |
|---|---|---|---|---|
| YAP1 | 1:200 | (A) | (63.7) sc-101199 | Santa Cruz |
| Vinculin | 1:200 | (A) | V9131 | Merck |
| Collagen 1A1 | 1:100 | (B) | 91144S | Cell Signaling |
| Collagen 3A1 | 1:100 | (B) | PA5-27828 | Thermo Fisher |
| CTGF | 1:100 | (B) | Ab5097 | Abcam |
| Fibronectin | 1:100 | (B) | F3648 | Merck |
| Laminin | 1:25 | (B) | SAB4200719 | Merck |
| Periostin | 1:100 | (B) | Sc-49480 | Santa Cruz |

(A) 2.5 % BSA, PBS, 2h (RT) – O.N. (4 °C)

(B) 2.5 % BSA, PBS, O.N. (4 °C)

**Minimum Information Reporting in Bio−Nano Experimental Literature (*86*). Supplementary Table 2. Material characterization**

| Question | Yes | No |
|---|---|---|
| 1.1 Are "**best reporting practices**" **available** for the nanomaterial used? | X | |
| 1.2 If they are available, **are they used**? If not available, ignore this question and proceed to the next one. | | |
| 1.3 Are extensive and clear instructions reported detailing all steps of **synthesis** and the resulting **composition** of the nanomaterial? | X | |
| 1.4 Is the **size** (or **dimensions**, if non-spherical) and **shape of** the nanomaterial reported? | X | |
| 1.5 Is the **size dispersity** or **aggregation** of the nanomaterial reported? | X | |
| 1.6 Is the **zeta potential** of the nanomaterial reported? | X | |
| 1.7 Is the **density (mass/volume)** of the nanomaterial reported? | | X |
| 1.8 Is the amount of any **drug loaded** reported? 'Drug' here broadly refers to functional cargos (e.g., proteins, small molecules, nucleic acids). | | X |
| 1.9 Is the **targeting performance** of the nanomaterial reported, including **amount** of ligand bound to the nanomaterial if the material has been functionalised through addition of targeting ligands? | | X |
| 1.10 Is the **label signal** per nanomaterial/particle reported? For example, fluorescence signal per Particle for fluorescently labelled nanomaterials. | X | |
| 1.11 If a material property not listed here is varied, has it been **quantified**? | | NA |
| 1.12 Were characterizations performed in a **fluid mimicking biological conditions**? | X | |
| 1.13 Are details of how these parameters were **measured/estimated** provided? | X | |
| Explanation for **No** (if needed): | | |

**Supplementary Table 3. Biological characterization**

| Question | Yes | No |
|---|---|---|
| 2.1 Are **cell seeding details**, including **number of cells plated, confluency at start of Experiment,** and **time between seeding and experiment** reported? | X | |
| 2.2 If a standardised cell line is used, are the **designation and source** provided? | X | |
| 2.3 Is the **passage number** (total number of times a cell culture has been subcultured) known and reported? | X | |
| 2.4 Is the last instance of **verification of cell line** reported? If no verification has been performed, is the time passed and passage number since acquisition from trusted source (e.g., ATCC or ECACC) reported? | | X |
| 2.5 Are the results from **mycoplasma testing** of cell cultures reported? | | X |
| 2.6 Is the **background signal of cells/tissue** reported? (E.g., the fluorescence signal of cells without particles in the case of a flow cytometry experiment.) | X | |

| | | |
|---|---|---|
| 2.7 Are **toxicity studies** provided to demonstrate that the material has the expected toxicity, and that the experimental protocol followed does not? | X | |
| 2.8 Are details of media preparation (**type of media, serum,** any **added antibiotics**) provided? | X | |
| 2.9 Is a **justification of the biological model** used provided? | X | |
| 2.10 Is characterization of the **biological fluid** (*ex vivo*/*in vitro*) reported? For example, when investigating protein adsorption onto nanoparticles dispersed in blood serum, pertinent aspects of the blood serum should be characterised (e.g., protein concentrations and differences between donors used in study). | X | |
| 2.11 For **animal experiments**, are the ARRIVE guidelines followed? | | NA |
| Explanation for **No** (if needed): | | |

**Supplementary Table 4. Experimental details**

| **Question** | **Yes** | **No** |
|---|---|---|
| 3.1 For cell culture experiments: are **cell culture dimensions** including **type of well**, **volume of added media**, reported? Are cell types (i.e.; adherent vs suspension) and **orientation** (if non- standard) reported? | X | |
| 3.2 Is the **dose of material administered** reported? This is typically provided in nanomaterial mass, volume, number, or surface area added. Is sufficient information reported so that regardless of which one is provided, the other dosage metrics can be calculated (i.e. using the dimensions and density of the nanomaterial)? | X | |
| 3.3 For each type of imaging performed, are details of how **imaging** was performed provided, including details of **shielding, non-uniform image processing**, and any **contrast agents** added? | X | |
| 3.4 Are details of how the dose was administered provided, including **method of administration, injection location, rate of administration,** and details of **multiple injections**? | X | |
| 3.5 Is the methodology used to **equalise dosage** provided? | X | |
| 3.6 Is the **delivered dose** to tissues and/or organs (in vivo) reported, as % injected dose per gram of tissue (%ID $g^{-1}$)? | | NA |
| 3.7 Is **mass of each organ/tissue measured** and **mass of material** reported? | | NA |
| 3.8 Are the **signals of cells/tissues with nanomaterials** reported? For instance, for fluorescently labelled nanoparticles, the total number of particles per cell or the fluorescence intensity of particles + cells, at each assessed timepoint. | X | |

| | |
|---|---|
| 3.9 Are **data analysis details**, including **code used** for analysis provided? | X |
| 3.10 Is the **raw data** or **distribution of values** underlying the reported results provided? | X |
| Explanation for **No** (if needed): | |
